# HSP70-binding motifs function as protein quality control degrons

**DOI:** 10.1101/2021.12.22.473789

**Authors:** Amanda B. Abildgaard, Vasileios Voutsinos, Søren D. Petersen, Fia B. Larsen, Caroline Kampmeyer, Kristoffer E. Johansson, Amelie Stein, Tommer Ravid, Claes Andréasson, Michael K. Jensen, Kresten Lindorff-Larsen, Rasmus Hartmann-Petersen

**Affiliations:** The Linderstrøm-Lang Centre for Protein Science, Department of Biology, University of Copenhagen, Copenhagen, Denmark; Novo Nordisk Foundation Center for Biosustainability, Technical University of Denmark, Kgs. Lyngby, Denmark; Department of Biological Chemistry, The Alexander Silberman Institute of Life Sciences, The Hebrew University of Jerusalem, Jerusalem, Israel; Department of Molecular Biosciences, The Wenner-Gren Institute, Stockholm University, Stockholm, Sweden

**Keywords:** protein unfolding, protein degradation, proteasome, protein stability, protein quality control, chaperone

## Abstract

Protein quality control (PQC) degrons are short protein segments that target misfolded proteins for proteasomal degradation, and thus protect cells against the accumulation of potentially toxic non-native proteins. Studies have shown that PQC degrons are hydrophobic and rarely contain negatively charged residues, features which are shared with chaperone-binding regions. Here we explore the notion that chaperone-binding regions may function as PQC degrons. When directly tested, we found that a canonical Hsp70-binding motif (the APPY peptide) functioned as a dose-dependent PQC degron both in yeast and in human cells. In yeast, Hsp70, Hsp110, Fes1, and the E3 Ubr1 target the APPY degron. Screening revealed that the sequence space within the chaperone-binding region of APPY that is compatible with degron function is vast. We find that the number of exposed Hsp70-binding sites in the yeast proteome correlates with a reduced protein abundance and half-life. Our results suggest that when protein folding fails, chaperone-binding sites may operate as PQC degrons, and PQC-linked degradation therefore overlaps in specificity with chaperone binding. This sheds new light on how the PQC system has evolved to exploit the intrinsic capacity of chaperones to recognize misfolded proteins, thereby placing them at the nexus of protein folding and degradation.

**Significance Statement:** It is broadly accepted that misfolded proteins are often rapidly degraded by the ubiquitin-proteasome system, but how cells specifically recognize this immensely diverse group of proteins is largely unknown. Here we show that upon uncoupling of protein folding from protein degradation, a canonical chaperone-binding motif doubles as a degradation signal (degron), and that within the context of a Hsp70-binding region, many sequences are compatible with degron function. We find that degradation is correlated with the number of Hsp70-binding sites within a protein, and that the number of exposed Hsp70-binding sites in the yeast proteome correlates with more rapid degradation.

## Introduction

Proteins are diverse and dynamic macromolecules that typically fold into native structures that determine their individual functions. Nevertheless, proteins are at continuous risk of misfolding, which may be accelerated due to mutations, aging or environmental stress conditions. Misfolded proteins are typically prone to form aggregates and often display proteotoxicity via poorly understood mechanisms (Bucciantini et al., 2002). For example, several neurodegenerative disorders have been connected with the accumulation of toxic misfolded protein species (Bates et al., 2015; Poewe et al., 2017). Thus, securing that the proteome is folded and functional is indispensable for cellular fitness. The outcome of protein folding is monitored closely by the cellular protein quality control (PQC) system that relies on both chaperones and the ubiquitin-proteasome system (UPS). Molecular chaperones prevent protein aggregation by promoting *de novo* folding of newly synthesized proteins, refolding of misfolded proteins, and the assembly of protein complexes (Balchin et al., 2020, 2016; Dahiya and Buchner, 2019). Regulated by Hsp40 J-domain co-chaperones and nucleotide exchange factors (NEFs), the heat shock protein 70 (Hsp70) chaperones catalyze protein folding through ATP-dependent cycles of substrate binding and release (Kohler and Andréasson, 2020; Rosenzweig et al., 2019). The preferred binding site for Hsp70-type chaperones is a short linear stretch of hydrophobic residues flanked by positive charges (Rüdiger et al., 1997), and Hsp70 therefore does not appear to recognize specific sequence motifs, but rather the overall physical characteristics such as exposed hydrophobicity. As such regions are widespread in the proteome (estimated to be present once for every 36 residues) (Rüdiger et al., 1997) and typically buried within the core of folded proteins, chaperones may essentially engage any misfolded protein that exposes such a region. Hsp70 promotes folding by clamping onto these regions (Houben et al., 2020; Rousseau et al., 2006; Rüdiger et al., 1997), and if the regions are buried upon release, the target has become folded. However, if folding fails, the persistent misfolded proteins are targeted for degradation through various PQC pathways, including the ubiquitin-proteasome system (UPS). To facilitate degradation by the 26S proteasome, proteins are covalently linked to ubiquitin moieties through the sequential actions of E1, E2 and E3 enzymes (Dikic, 2017; Kwon and Ciechanover, 2017; Pohl and Dikic, 2019).

The pathway for PQC degradation of misfolded proteins is not fully resolved, and a key unsettled question is how the specificity is achieved, i.e. how does the PQC system distinguish misfolded proteins from the many proteins that should not be degraded. Typically, the UPS obtains specificity through the substrate preference of E3 ubiquitin-protein ligases. Accordingly, a handful of E3s have been connected with PQC-linked degradation of misfolded proteins in the cytosol and nucleus (Hickey et al., 2021; Oh et al., 2018; Samant et al., 2018), but the exact substrate preferences of the PQC E3s are largely unknown. In general, E3s recognize regions in target proteins termed degradation signals or *degrons* (Varshavsky, 1991). The region via which the protein is interacting with the E3, and thus is determining the specificity, is called the primary degron, but in the following referred to as *degron* (Guharoy et al., 2016b, 2016a).

Unlike the degrons involved in the regulated degradation of folded proteins, such as the KEN-box, the N-end and C-end degrons (Bachmair et al., 1986; Koren et al., 2018; Pfleger and Kirschner, 2000; Timms and Koren, 2020; Varshavsky, 2011), the PQC degrons are not well described. However, a few examples such as *Deg1* and *CL1* have been characterized in detail. The *Deg1* degron is situated in the a1-interaction site of the yeast α2 protein, while *CL1* is an artificial 16-residue sequence. Both depend on hydrophobic residues mediating the degradation by the E3 Doa10 (Gilon et al., 1998; Johnson et al., 1998; Ravid et al., 2006). Recent studies show that in yeast, PQC degrons in general contain stretches of hydrophobic residues and are depleted for negatively charged residues (Johansson et al., 2022; Mashahreh et al., 2022), and similar amino acid preferences have been observed in mammalian cells (Koren et al., 2018). As these features are shared with chaperone-binding regions (Houben et al., 2020; Koopman and Rüdiger, 2020; Rousseau et al., 2006), this suggests that chaperones might actively engage in substrate selection for PQC-linked degradation. Such a mechanism is also supported by data on the PQC-linked E3s, Doa10 and Ubr1, which both collaborate with molecular chaperones for degradation via poorly understood mechanisms (Amm et al., 2015; Heck et al., 2010; Kriegenburg et al., 2014; Maurer et al., 2016; Shiber et al., 2013; Singh et al., 2020; Summers et al., 2013). For Ubr1, a direct interaction with Hsp70 has been observed (Summers et al., 2013), indicating that some PQC E3s may delegate substrate recognition to the chaperones. Similarly, the human CHIP E3 directly interacts with Hsp70 and Hsp90 chaperones to ubiquitylate bound protein substrates (Höhfeld and Jentsch, 1997), while the chaperone NEFs BAG1 and Hsp110 mediate the release of bound substrates at the 26S proteasome (Abildgaard et al., 2020; Gersing et al., 2021; Höhfeld and Jentsch, 1997; Kandasamy and Andréasson, 2018; Kriegenburg et al., 2014). The broad substrate specificity of Hsp70 and its involvement in protein degradation suggest that it plays a fundamental and direct role in targeting misfolded proteins to the UPS via binding their short PQC degrons. Here, we directly test if Hsp70-binding regions can function as degrons in cytosolic quality control (CytoQC). We found that the canonical Hsp70-binding motif RLLL, embedded within the so-called APPY peptide (Montgomery et al., 1999; Schmid et al., 1994, 1992), shows an inherent degron capacity in both yeast and human cells. We show that the sequence context surrounding the RLLL motif is critical for its degron function, while a broad sequence space within the RLLL motif is compatible with degron activity. In conclusion, our data indicate that many cytosolic PQC degrons share features with regions adept at engaging the Hsp70 substrate-binding cleft.

## Results

### The consensus Hsp70-binding motif (RLLL) functions as a degron

To test the notion that chaperone-binding sites may function as PQC degrons, we first implemented a growth-based degron assay based on a Ura3-HA-GFP fusion as a reporter expressed in yeast cells (Geffen et al., 2016). In this system, the degron capacity of a C-terminally attached peptide can be monitored as a growth phenotype due to the degradation of the Ura3 reporter protein (Fig. 1A). As an example, fusion to the 16-residue *CL1* degron sequence (ACKNWFSSLSHFVIHL) (Gilon et al., 1998) promotes proteasomal degradation of the reporter, which results in growth deficiency on uracil-depleted medium (Fig. 1B). The growth defect is due to low cellular abundance of the reporter, as shown by Western blotting (Fig. 1C) and since treating cells with the proteasomal inhibitor, bortezomib (BZ), restores protein levels, which in turn and rescues the growth defect (Fig. 1BC). Moreover, as the Ura3 enzyme converts 5-fluoroorotic acid (5-FOA) to the toxic 5-fluorouracil compound (Boeke et al., 1984), degradation of the reporter provides cells with resistance to 5-FOA (Fig. 1B) (Gilon et al., 1998).

**Figure 1.**
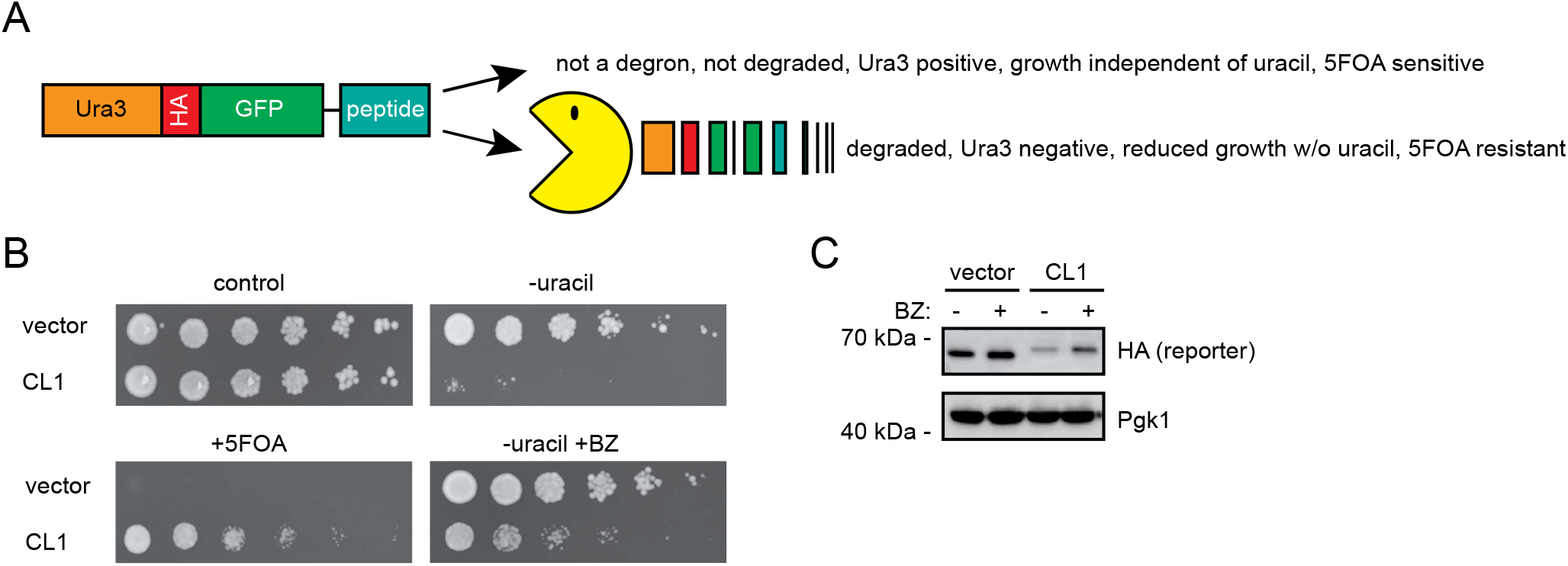
Assessing degron activity with the Ura3-HA-GFP reporter system. (A) A schematic representation of the Ura3-HA-GFP reporter system. In-frame fusion of degrons to the C-terminus of the reporter will lead to proteasomal degradation (Pac-Man) leading to a 5-FOA resistant phenotype. (B) As a proof of principle for the reporter system, the growth of wild-type cells expressing the reporter vector with and without fusion to the CL1 degron was compared by dilutions on solid media. Note that the growth defect of the CL1 strain on media without uracil is suppressed by the addition of the proteasome inhibitor bortezomib (BZ), while in the presence of 5-FOA, the CL1 strain is able to form colonies. (C) Analysis of protein levels in vector or CL1 cells after 4 hours of bortezomib (BZ) treatment by SDS-PAGE and Western blotting, using antibodies to the HA-tag in the reporter. Blotting against Pgk1 served as loading control.

Using this system, we continued to test the so-called APPY peptide, which we term here as 1xRLLL. This is a canonical Hsp70-binding 22-mer peptide (CALLQS**RLLL**SAPRRAAATARY), originally identified in the precursor of chicken mitochondrial aspartate aminotransferase (Montgomery et al., 1999; Schmid et al., 1994, 1992). The central RLLL motif is critical for binding Hsp70, and the structure of Hsp70 bound to an RLLL peptide (Zhu et al., 1996) shows the RLLL sequence placed within the Hsp70 substrate binding groove (Fig. 2A). We prepared a set of reporter-peptide fusions to test the degron capacity of the RLLL motif (Fig. 2B). To potentially increase avidity, we introduced the entire 22-mer APPY peptide in one (1xRLLL), two (2xRLLL) or three (3xRLLL) copies in the C-terminus of the Ura3-HA-GFP reporter. We also constructed variants, RAAA and DAAA, in the central RLLL motif in APPY. These are expected to have decreased Hsp70-binding capacity and are thus predicted to confer intermediate or poor degron capacities, respectively (Rüdiger et al., 1997). The reporter-constructs were expressed in yeast cells, and protein levels were analyzed by microscopy (Fig. 2C) and Western blotting (Fig. 2D). The RLLL-fused reporter was found at low levels, which decreased further with increasing repeats of the APPY peptide (2xRLLL and 3xRLLL). In contrast, for the RAAA and DAAA variants, the levels were, respectively, partly or completely restored, thus reflecting their predicted Hsp70 affinities from *in vitro* studies (Rüdiger et al., 1997). This indicates that degradation of the reporter correlates with the Hsp70-binding ability. Treating cells with the translational inhibitor, cycloheximide (CHX), caused the level of the RLLL-fused reporter protein to decrease further, while inhibition of the proteasome caused an increase (Fig. 2E), proving that the reduced protein level is caused by proteasomal degradation. Meanwhile, the levels of the DAAA-variant of APPY remained unchanged by both treatments. When we then assessed the growth phenotypes of the reporter-constructs on uracil-depleted medium, we observed that proteasome inhibition rescued the growth of cells with degron-fused reporters (Fig. 2F). Collectively, these results show that decreased cellular abundances are due to proteasomal degradation of the RLLL-linked reporters, and that repeats of the degron is more efficient than a single copy.

**Figure 2.**
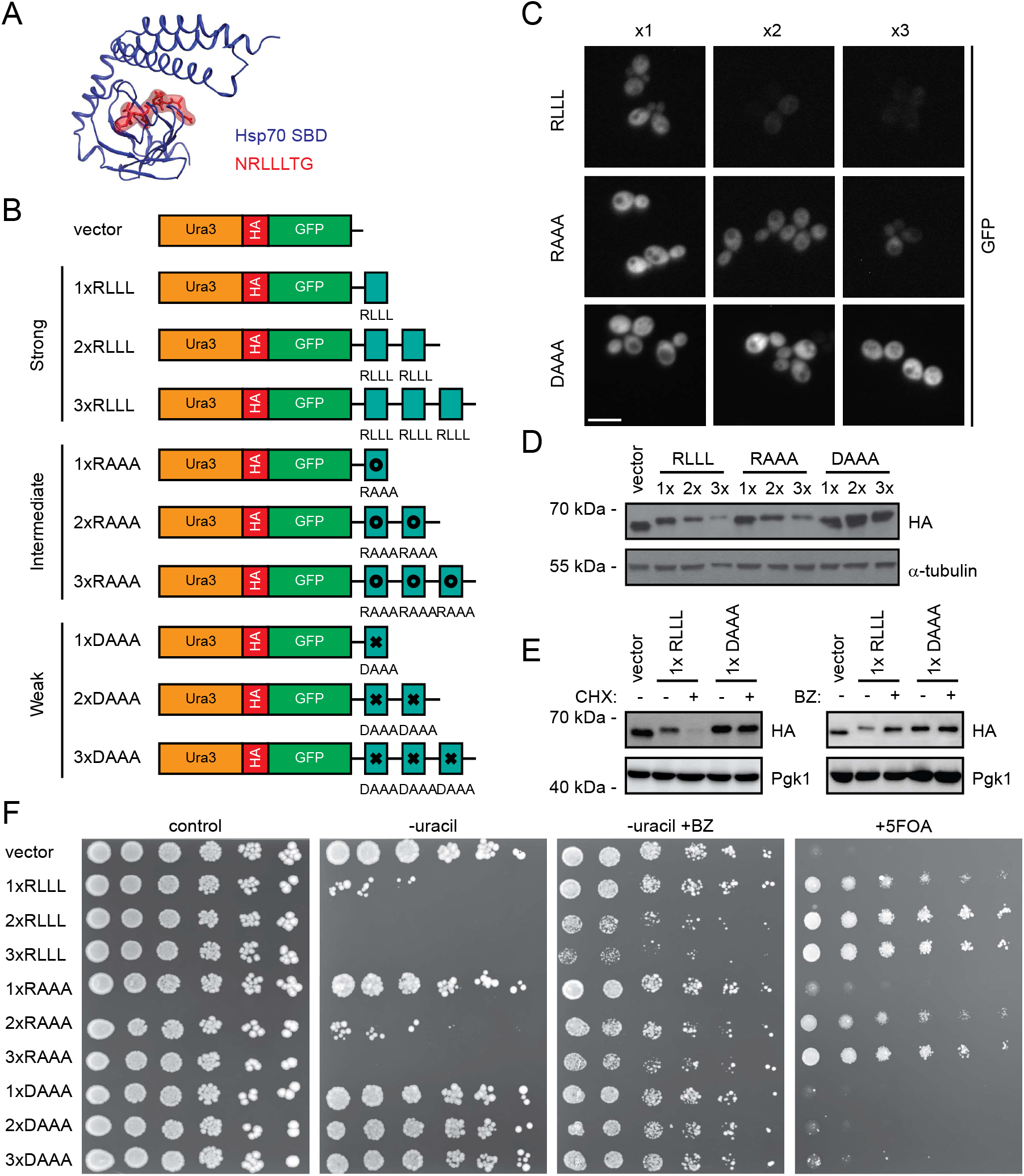
Investigation of the degron capacity of the canonical Hsp70-binding motif, RLLL. (A) Structure of the Hsp70 substrate-binding domain (SBD) (blue) in complex with the Hsp70-binding peptide, NRLLLTG (red) (PDB: 4PO2) (Zhang et al., 2014). (B) Illustration of APPY-variants fused in one, two or three copies to the Ura3-HA-GFP-reporter. Note that the entire 22-residue APPY peptide was used. From the literature (Rüdiger et al., 1997), we predicted strong (RLLL), intermediate (RAAA) and weak (DAAA) degron motifs. (C) Cells carrying the indicated fusions were analyzed by microscopy for GFP fluorescence. Bar, 10 μm. (D) Cell lysates of cells carrying the constructs listed in panel (B) were prepared and compared by SDS-PAGE and Western blotting using antibodies to the HA-tag. Tubulin served as loading control. (E) Analysis of the indicated protein levels by SDS-PAGE and Western blotting from cells treated with the translation inhibitor cycloheximide (CHX) (left panel) or the proteasome inhibitor bortezomib (BZ) (right panel). Pgk1 served as loading control. (F) Growth assay of cells carrying the constructs listed in panel (B).

### The RLLL-degron relies on chaperones, co-chaperones, and the E3 Ubr1

To further explore the APPY RLLL-degron, we next analyzed the solubility of the RLLL fusion constructs, since previous studies have shown a positive correlation between protein insolubility and degradation of PQC targets (Abildgaard et al., 2019; Fredrickson et al., 2013; Gallagher et al., 2014; Jones et al., 2020; Summers et al., 2013). To test this for the RLLL peptides, we separated whole cell lysates into soluble and insoluble fractions by centrifugation. This revealed that the RLLL motif induces insolubility of the reporter construct (Fig. 3A). While the solubility of the reporter decreased with the number of RLLL motifs in the construct, mutations within the RLLL motif reduced (RAAA) or eliminated (DAAA) this effect (Fig. 3A). Thus, the degron-activity of the RLLL variants correlates with protein insolubility.

**Figure 3.**
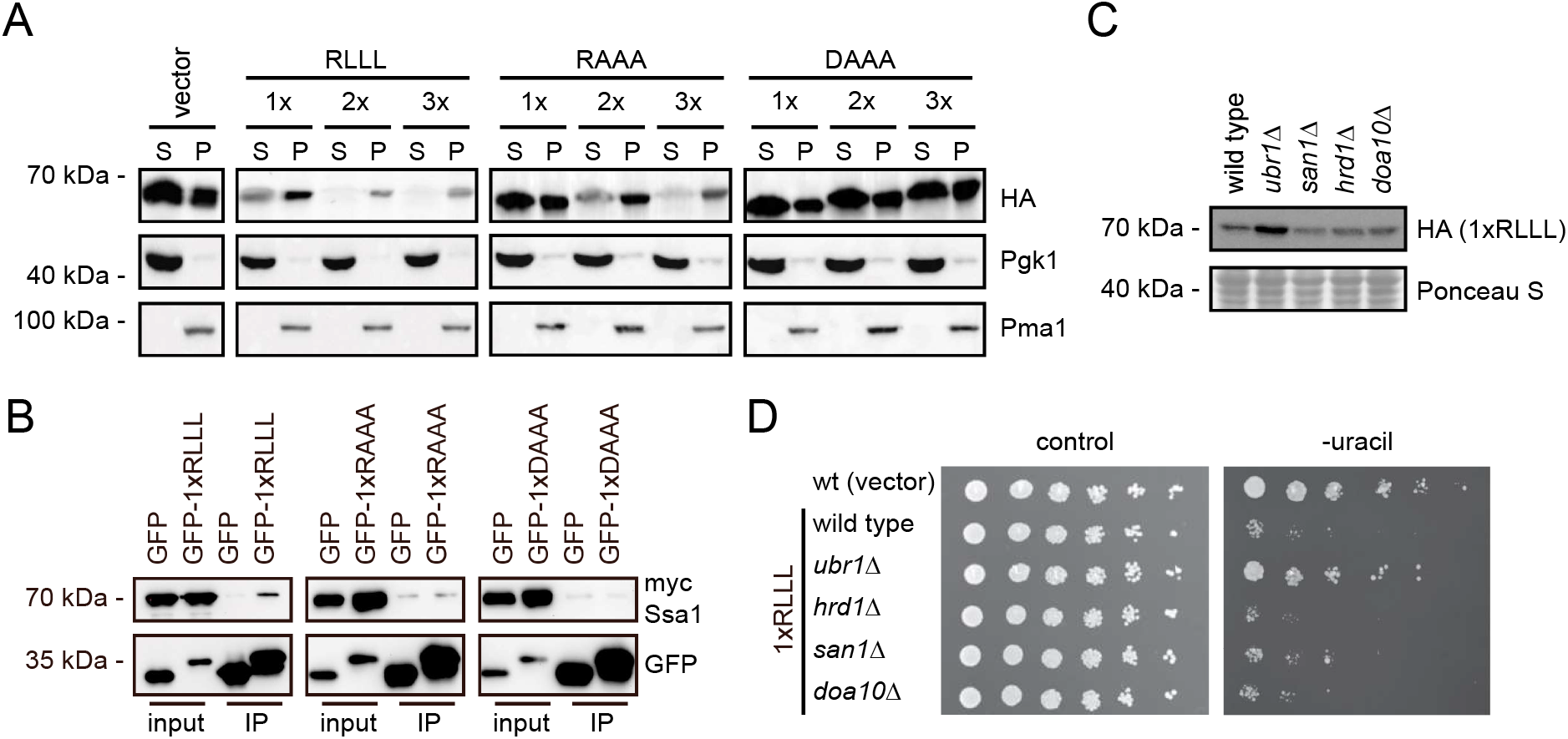
Characterization of the RLLL degron. (A) The solubility of the reporter proteins was analyzed by fractionating cell extracts into soluble supernatant (S) and insoluble pellet (P) fractions by centrifugation. The protein levels were analyzed by SDS-PAGE and Western blotting using antibodies to the HA-tag in the reporter. Pgk1 served as a loading control for the soluble protein fractions, while Pma1 served as loading control for the insoluble protein fractions. (B) Myc-tagged Ssa1 (Hsp70) was co-immunoprecipitated (co-IP) with the indicated GFP fusions using GFP-trap resin from cultures treated with bortezomib. ATP was not added to the buffers used for the co-IPs. The precipitated protein was analyzed by SDS-PAGE and blotting with antibodies to myc and GFP. (C) The protein levels of the RLLL degron were compared in the indicated yeast strains by SDS-PAGE and blotting for the HA-tag on the reporter. A Ponceau S staining of the membrane is included as loading control. (D) The dependence of E3 ligases for targeting the RLLL degron was analyzed by growth assays on solid media using the indicated null mutants. Wild-type cells transformed with the reporter vector alone were included for comparison.

Next, we tested if we could detect interaction with Hsp70 using a high copy expression vector for the GFP-1xRLLL fusion in immunoprecipitation assays with myc-tagged yeast Hsp70 homologue Ssa1. Since the 2xRLLL and 3xRLLL fusions were insoluble, we could not test for Hsp70 interactions with these constructs. To limit the degradation of the degron fusions, BZ was added to the cultures 4 hours prior to lysis. As anticipated, the GFP-1xRLLL fusion protein interacted with Ssa1, while the GFP control did not (Fig. 3B). We noted that only a fraction of the total Ssa1 was precipitated. Presumably, this is linked to the transient nature of this enzyme-substrate complex. However, the interaction seems specific, since replacing the RLLL motif in the APPY peptide with either RAAA or DAAA eliminated binding. Accordingly, the level of the reporter-RLLL degron fusion increased when Hsp70 was inhibited with myricetin (Chang et al., 2011) (Fig. S1A) or with YM01 in human cells (see below). By growth assays and western blotting, we explored the potential involvement of a number of co-chaperones and E3 ligases. In yeast, Hsp70-dependent protein degradation depends on the Hsp70 NEFs Fes1 (Gowda et al., 2018, 2013) and the Hsp110 orthologues Sse1 and Sse2 (Andréasson et al., 2008), which mediate substrate release and degradation at the 26S proteasome (Kandasamy and Andréasson, 2018). Accordingly, the level of the RLLL fusion was increased in *fes1*Δ cells (Fig. S1B) and in the *sse1-200sse2*Δ (Kaimal et al., 2017) temperature-sensitive double-mutant strain (Fig. S1C), indicating that these NEFs are required for the degradation of the RLLL degron. We also analyzed strains deleted for various Hsp40 J-domain co-chaperones (Supplementary Fig. S1BD). We found that deletion of the J-domain proteins Jjj3, Apj1, Zuo1 and Caj1 increased cellular abundance of the RLLL-fused reporter, suggesting that they are directly or indirectly (see Discussion) involved in the degradation of RLLL. Intriguingly, in strains lacking the yeast-specific Hsp104 and Jjj2 (Supplementary Fig. S1BD), a reduced growth on uracil-depleted medium and decreased steady-state levels were observed. This suggests that these components, directly or indirectly, block RLLL-degradation, perhaps by affecting the aggregation or solubility of the RLLL reporter.

Finally, we tested selected PQC E3 ubiquitin-protein ligases and observed a strong effect in the *ubr1*Δ strain (Fig. 3CD), indicating that Ubr1 plays a role in degradation of the RLLL reporter, which is in line with its previously published role in PQC (Summers et al., 2013). Accordingly, and as reported before (Summers et al., 2013), we observed an interaction between Ubr1 and Ssa1 by co-precipitation (Supplementary Fig. S1E).

### Possible degron sequences within the context of the APPY peptide

Having established that the APPY peptide functions as a PQC degron, we next analyzed the sequence properties connected to its degron activity. First, we tested the role of residues flanking the RLLL sequence by mutagenesis. Both growth assays (Fig. 4A) and protein abundances (Fig. 4B) of mutant reporter-constructs showed that the introduction of negative charges near the RLLL sequence disrupts its degron capacity. This may also indicate that the degron function relies on binding to Hsp70, which is known to be highly sensitive to the presence of negative charges flanking the binding site (Rousseau et al., 2006). As a further control to support that the RLLL fragment is likely exposed and the mutations do not induce structure, a circular dichroism (CD) analysis of selected peptide variants confirmed that the peptides hold no regular secondary structure (Supplementary Fig. S2).

**Figure 4.**
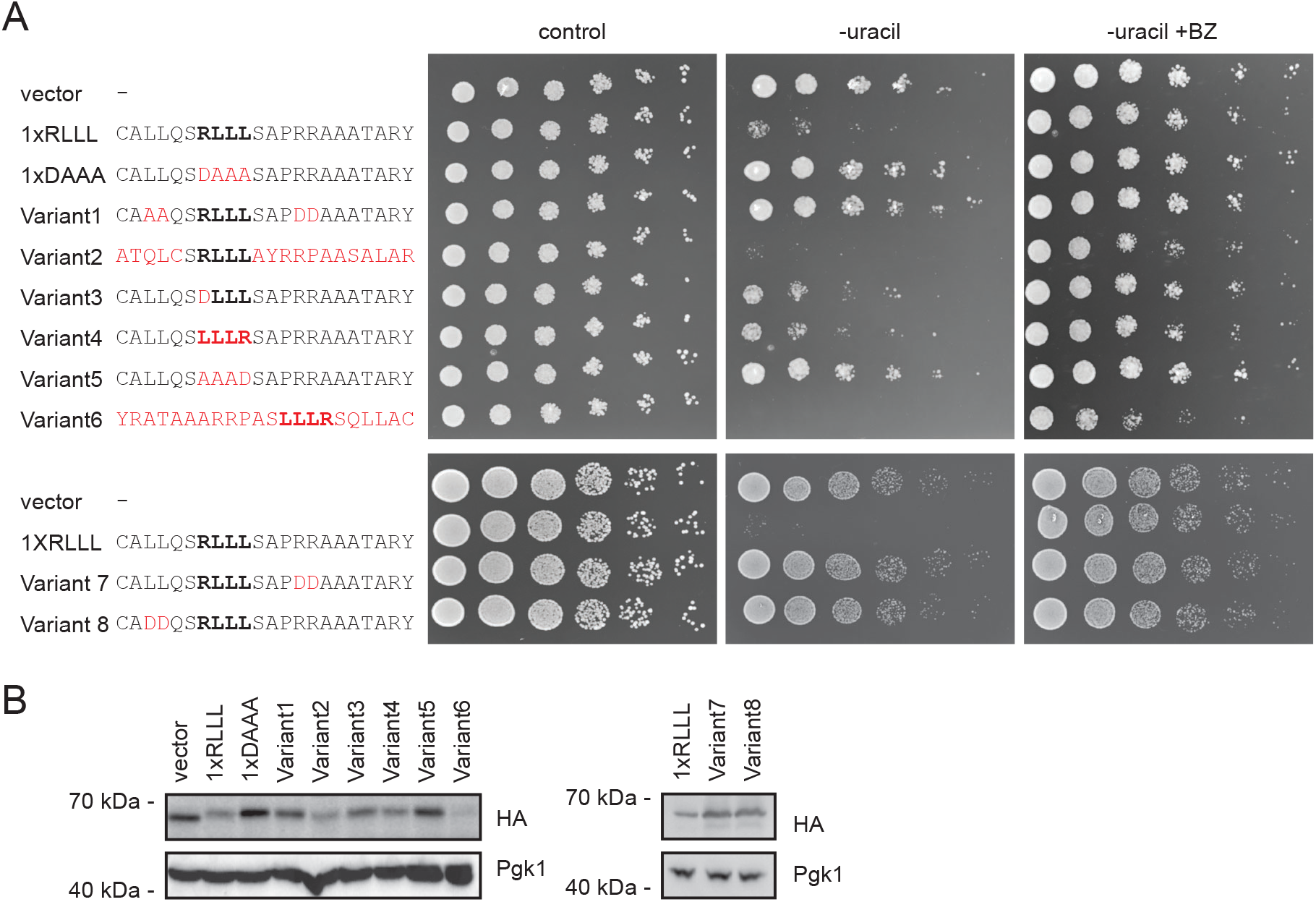
The RLLL degron sequence-dependence. (A) Fusion of the Ura3-HA-GFP-reporter to the listed APPY variants induced growth phenotypes that were assessed on solid media. The RLLL motif is highlighted (bold) and amino acid substitutions are shown in red. (B) The abundance of variants from panel A was compared by SDS-PAGE and Western blotting of whole-cell lysates using antibodies to HA. Pgk1 served as a loading control.

Next, we aimed at exploring the importance of the central RLLL sequence—in context of the APPY peptide—for its degron function. To that end, we constructed a library by randomization of the four amino acid positions in the RLLL motif of the APPY peptide downstream of the Ura3-HA-GFP reporter. To maximize the diversity of the library, randomization was achieved by insertion of four degenerate codon sets in one of the primers used for PCR prior to assembly by *in vivo* gap repair (Fig. 5A). Transformed cells were collected and plated on 5-FOA plates (Fig. 5A). By comparing the number of plated colonies from the treated and untreated libraries, we found that <0.5% of the library displayed resistance to 5-FOA and can thus be assumed to hold degron capacity. This indicates that the library is not skewed towards sequences with degrons. Illumina sequencing across the 3’ end of the *URA3-HA-GFP* reporter revealed large sequence variation among the possible degron motifs (Supplementary File 1), suggesting that either: 1) many specific degron sequences are targeted in parallel by multiple E3s, and/or 2) many different degron sequences are targeted by a mechanism of broad specificity. After quality control of our sequencing data, we found ca. 18,000 of the 160,000 possible tetrapeptide sequences in the input library, yet only 883 sequences were recovered after selection on 5-FOA. Analysis of the degron sequences revealed a generally high flexibility within the sequence space of the degron motifs. We found an overall enrichment of hydrophobic residues (Fig. 5B), while polar residues were generally depleted. For example, Leu, Met, Trp, Tyr, Phe, and Ile are all enriched (to various extents) and Gln, Thr, Asp, Glu and His are all depleted. There are, however, some exceptions such as an enrichment of Asn and Gly and a depletion of Ala. Thus, while the degrons are overall hydrophobic, the signal appears more complex than just hydrophobicity. All identified degrons are listed in the supplemental material (Supplementary File 1).

**Figure 5.**
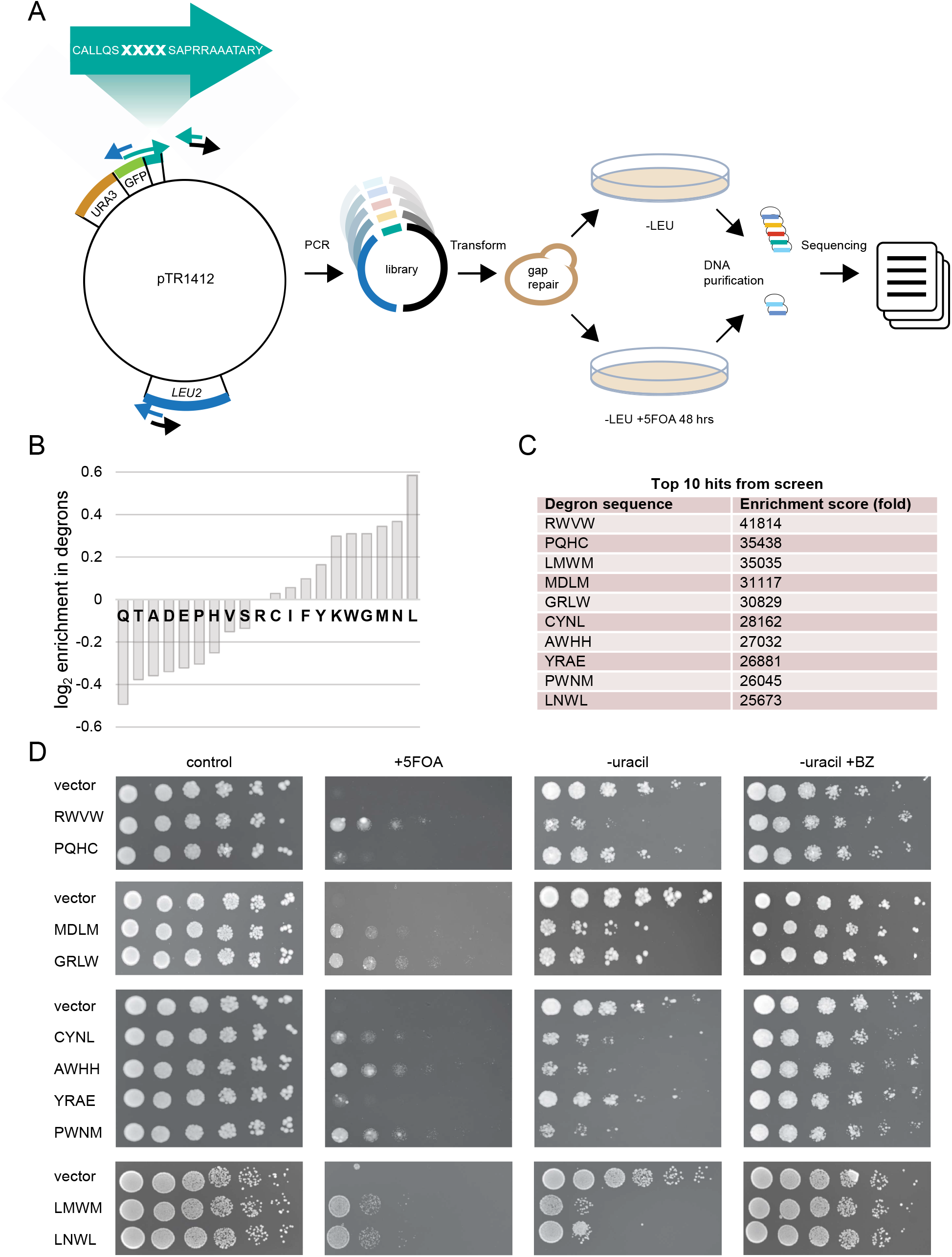
Screening reveals a large sequence space of functional degrons with the RLLL motif. (A) Schematic overview of the library construction and screen. Three primer pairs were designed to cover the pTR1412 vector, encoding the Ura3-HA-GFP-reporter. Four amino acid positions in the shown C-terminal peptide were randomized by using a primer carrying TriMix20 at the bold-faced positions marked X. PCR amplification yielded three partially overlapping fragments, which were assembled by gap repair following transformation into yeast. Transformants were selected on media without leucine (-LEU) and on media without leucine but with 5-FOA (-LEU+5-FOA) to select for degrons. Finally, DNA was extracted and sequenced. (B) Enrichment and depletion of amino acid types in degrons. The plot shows the log_2_ of the ratio of the fractions of each amino acid found in the top100 peptides relative to the full set of peptides seen in the screen. Residues with a positive value are enriched in degrons and those with negative values are depleted. (C) The ten most prevalent sequences in the 5-FOA-treated library are listed along with the enrichment scores. (D) The growth of the most prevalent sequences from the 5-FOA library was compared on solid media.

We note that it is possible that the cloning and screening strategy outlined above may result in some false positives, for example due to mutations in the *URA3* gene. We therefore selected the ten most prevalent sequences from the 5-FOA resistant library (Fig. 5C) and tested them individually. Their degron capacities were examined by analyzing resistance to 5-FOA as well as their growth on uracil-depleted medium with and without BZ (Fig. 5D). This confirmed strong degron capacities in eight motifs (RWVW, LMWM, MDLM, GRLW, CYNL, AWHH, PWNM, and LNWL) while two motifs (PQHC and YRAE) did not have significant degron activity. We conclude that many sequences may replace the central RLLL motif to retain the degron potential of APPY.

### Proteome-wide correlation of Hsp70-binding motifs and protein degradation

To examine the role of Hsp70 further we used Limbo, a predictor for peptide binding to *E. coli* Hsp70 (DnaK) (Durme et al., 2009), to predict binding of Hsp70 to all the variants of the APPY sequence that we screened. A high Limbo score indicates a greater likelihood of binding, and while we do not find an overall correlation between the sequencing counts and the prediction of the Hsp70 binding, we do find that those sequences most enriched after treatment with 5-FOA have a greater predicted likelihood for binding to Hsp70. For example, while the average Limbo score across all 18,706 sequences is 1.79±0.04 (mean ± standard error of the mean), the ten most prevalent sequences from the 5-FOA resistant library have an average Limbo score of 5±1 and the average score of the top-100 sequences is 3.0±0.5 (Fig. 6A). Thus, many potent degron sequences are predicted to bind to the substrate-binding site of Hsp70.

**Figure 6.**
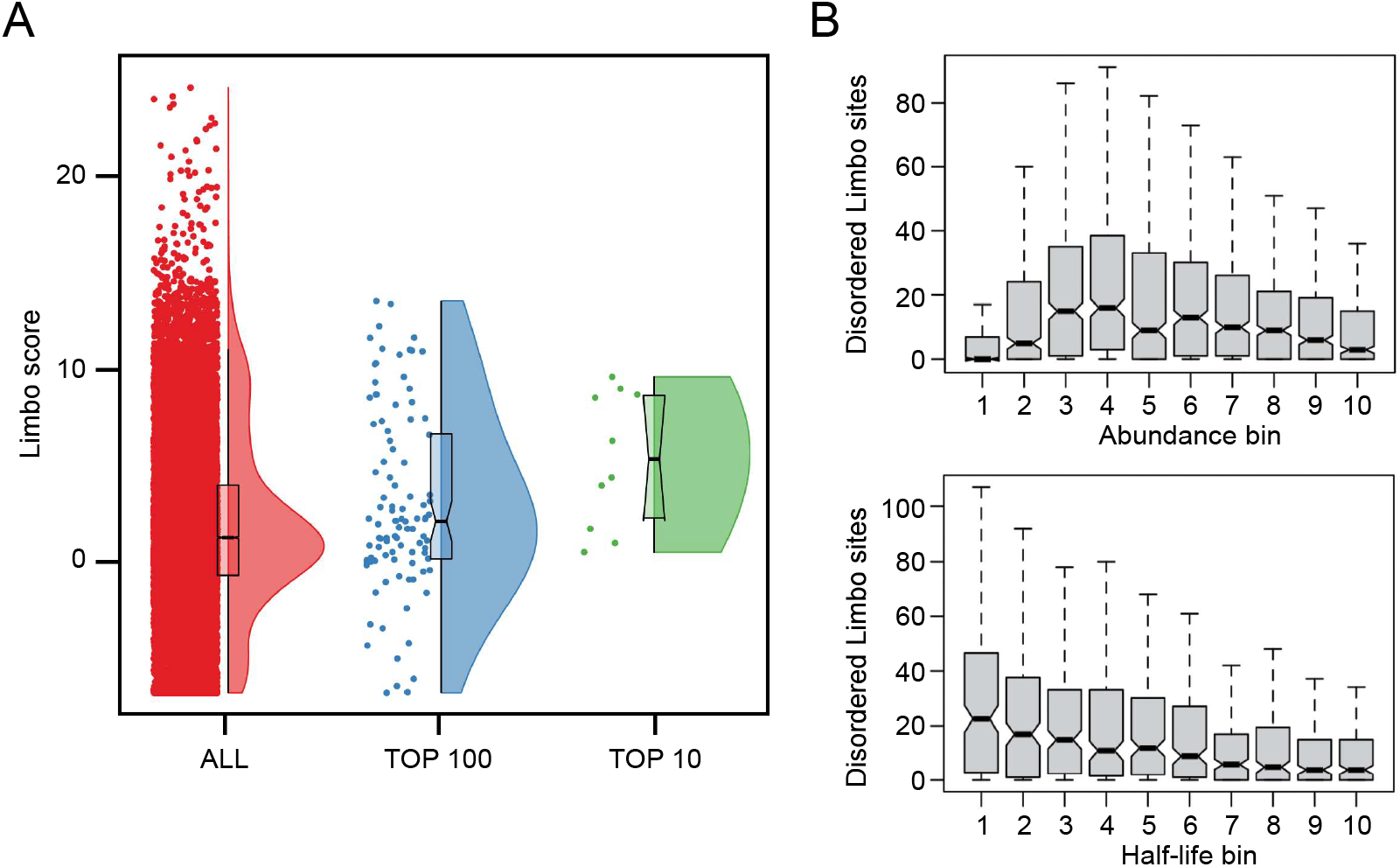
The number of exposed Hsp70-binding sites correlates with protein turnover. (A) The Limbo predictor for peptide binding to *E. coli* Hsp70 (DnaK) (Durme et al., 2009) was used to analyze all 18,706 sequences (red), or the top 100 sequences (blue) or the ten most prevalent sequences (green) from the 5-FOA resistant library. A high Limbo score indicates a greater likelihood for binding to Hsp70. (B) Proteome wide analysis of predicted Hsp70-binding sites. The number of residues that are both predicted to be Hsp70-binding sites and disordered, and thus solvent exposed, is different for proteins with different levels of abundance and half-life. Very low abundance proteins are an exception to this trend which has been suggested to reflect the absence of selection against exposed degrons when the protein concentrations are lower than the binding affinity of harmful interactions (Dubreuil et al., 2019).

Inspired by previous work (Macossay-Castillo et al., 2019), we then asked the question whether there is a correlation between predicted Hsp70 binding and protein abundance more generally. We thus calculate Limbo scores for the yeast proteome and consider only those residues predicted to be disordered, and thus solvent accessible. We find that the mean number of Hsp70-binding sites among these residues differs for different levels of abundance and half-life, indicating that exposed Hsp70-binding sites might influence protein abundance and half-life in general (Fig. 6B).

### The APPY peptide acts as a chaperone-dependent degron in human cells

Next, we tested if the APPY peptide also functions as a degron in human cells. To that end, we utilized a human HEK293T cell line modified to carry a “landing pad” for site-specific integration (Matreyek et al., 2020) (Fig. 7A). Upon integration of the expression plasmid at the landing pad, the BFP-iCasp9-Blast^R^ coding sequences are displaced and non-recombinant cells can be identified based on expression of BFP and selected against by adding AP1903 (Rimiducid) which induces apoptosis of iCasp9 positive cells (Fig. 7A). As the plasmid lacks a promoter, any non-recombined plasmids should not be expressed, while correct Bxb1-mediated site-specific integration in the landing pad locus should lead to expression of GFP-fused to the APPY peptide. Similar to a related system for analyzing degrons in human cells (Koren et al., 2018), the construct also allowed for correction of cell-to-cell variations in expression by encoding mCherry after an internal ribosomal entry site (IRES) (Fig. 7A). Thus, by flow cytometry it is possible to quantify the GFP-APPY abundance as the GFP:mCherry ratio. In agreement with our observations in yeast (Fig. 2CDF), adding the APPY peptide to the C-terminus of GFP in one, two and three copies led to reduced protein levels (Fig. 7B). Also, in agreement with the observations in yeast (Fig. 2EF), addition of the proteasome inhibitor bortezomib (BZ) led to an increase in the 1xRLLL APPY variant, whereas the level of 1xDAAA APPY was unaffected (Fig. 7C). To test if the abundance of the GFP-APPY variants depends on Hsp70, we compared the levels of the GFP-APPY variants in cells treated with the Hsp70 inhibitor YM01. The level of 1xRLLL APPY increased upon blocking Hsp70, whereas the 1xDAAA APPY variant was unaffected (Fig. 7D). Finally, as a further control for Hsp70 involvement, we used qPCR to test if expression of the 1xRLLL, 1xRAAA and 1xDAAA APPY variants resulted in an induction of Hsp70. This revealed about 3.5 fold induction of Hsp70 (HSPA1A and HSPA1B) in response to 1xRLLL APPY expression. The 1xRAAA APPY variant led to ~2.5 fold induction, while the effect of the 1xDAAA variant was insignificant (Fig. 7E). Based on these results we conclude that also in human cells the APPY peptide, and in particular its central RLLL motif, functions as a chaperonedependent degron.

**Figure 7.**
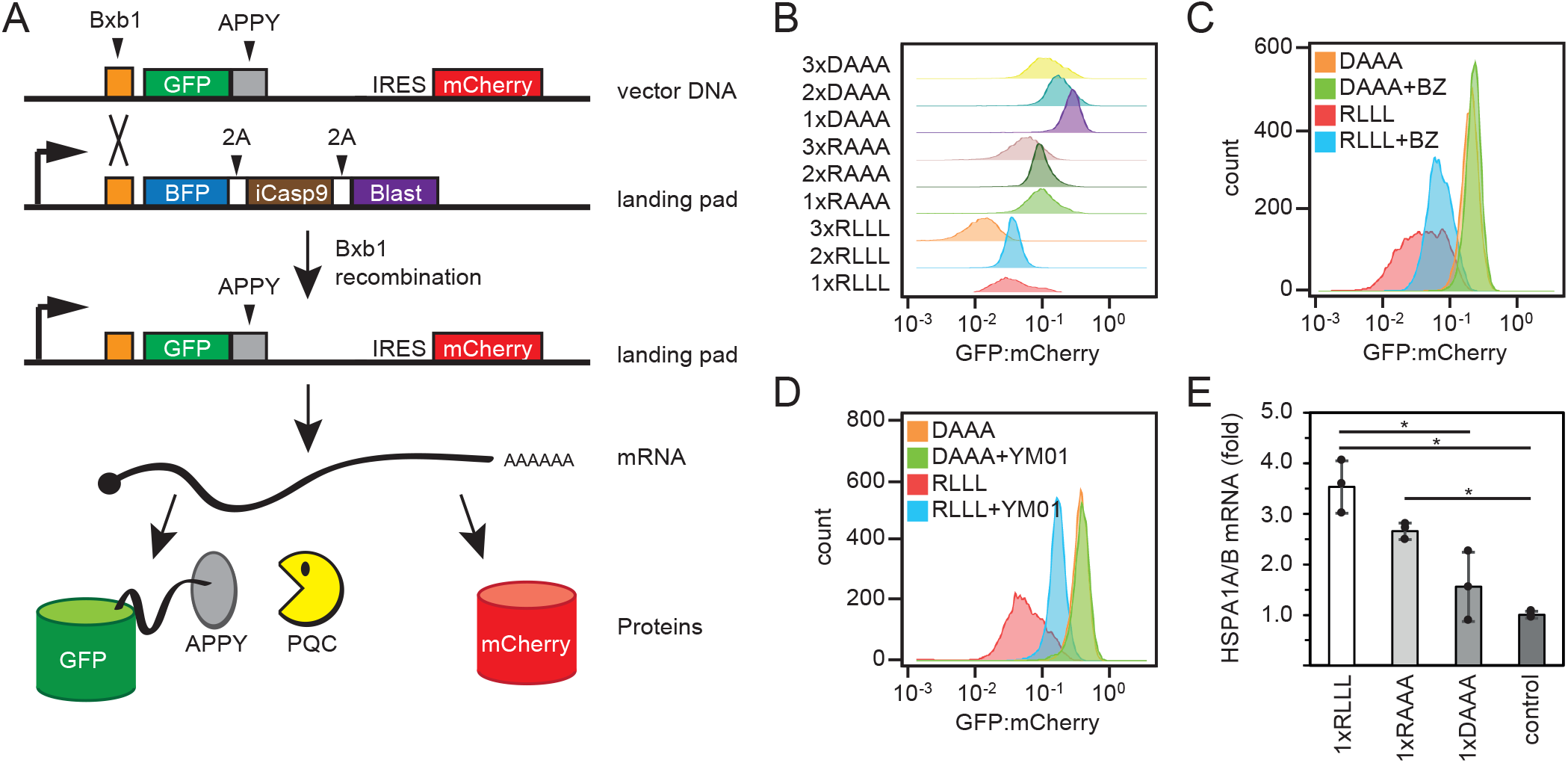
The APPY peptide functions as a degron in human cells. (A) Upon transfection of the HEK293T landing pad cells, BxbI catalyzes site-specific recombination between BxbI sites in the vector and the landing pad, leading to single-copy expression of GFP-APPY from the Tet-on promoter. As the vector also contains an internal ribosomal entry site (IRES) followed by mCherry, the GFP-APPY levels can be normalized to the mCherry levels. 2A signifies self-cleaving peptides. (B) GFP:mCherry ratio distributions of cells expressing the indicated GFP-APPY variants, obtained by flow-cytometry of ~10,000 cells. (C) GFP:mCherry ratio distributions of cells expressing the GFP-APPY (RLLL) peptide or the DAAA variant of APPY after 5 hours treatment with 15 μM bortezomib (BZ) or DMSO (solvent control), obtained by flow-cytometry of ~10,000 cells. (D) GFP:mCherry ratio distributions of cells expressing the APPY (RLLL) peptide or the DAAA variant of APPY after 22 hours treatment with 5 μM YM01 or no treatment, obtained by flow-cytometry of ~10,000 cells. (E) Relative expression of the *HSPA1A* and *HSPA1B* genes derived from qPCR. The results were normalized to the expression of non-recombinant cells. n=3, error bars show the standard deviation, asterisks indicate significant differences (p < 0.05) based on a two-tailed unpaired student’s t test.

## Discussion

In this study, we demonstrate that a region adept at engaging the substrate binding cleft of Hsp70 doubles as a PQC degron. Accordingly, Hsp70 binding sites likely play a role in both protein (re)folding and protein degradation. In our experimental systems, the APPY chaperone-binding motif is artificially grafted onto the Ura3-HA-GFP (yeast) and GFP (human) reporters, and since the chaperones cannot refold and bury these fragments, this effectively uncouples degradation from folding, thus allowing us to study the consequences of exposing these regions. Although this model system differs from PQC degron-exposure in a natural full-length misfolded protein, which while still reliant on Hsp70 is indisputably more complex (Kampmeyer et al., 2022), our findings lead us to suggest a model (Fig. 8). First, exposure of a chaperone-binding motif, induced by misfolding, leads to initial chaperone engagement. Then, in a second step, if refolding is not possible or slow, PQC E3s will be recruited to the chaperone-substrate complex and target the misfolded protein for proteolysis through ubiquitylation. Thus, if the substrate (re)folds after chaperone engagement, the chaperone-binding motif will become buried and will no longer target the protein for proteolysis. Proteins that persistently occupy misfolded conformations will, however, remain associated with Hsp70 and eventually be targeted for degradation. Importantly, PQC E3s may also directly interact with motifs that overlap with chaperone-binding regions and recruitment of E3s through direct chaperone-binding is therefore not strictly required. Regardless, according to this model, any substrate that Hsp70 binds will risk degradation. This provides a new perspective on how Hsp70 chaperones perform triage of bound substrates. A prediction from this model is that the refolding rates of substrates are more rapid than association with components required for PQC-linked degradation. According to the *Saccharomyces* Genome Database, the combined abundance of Ssa1 and Ssa2 is ~500,000 molecules/cell, while the PQC E3s are all present at levels roughly two orders of magnitude lower. Hence, a simple stochastic model where PQC E3s randomly ubiquitylate chaperone clients may result in specific clearance of irreversibly misfolded proteins or proteins with exposed chaperone binding sites. An additional level of specificity and regulation can then be achieved through repeated cycles of binding and release by the chaperone, which involve recruitment of specific co-chaperones, including J-domain proteins, NEFs like Hsp110 or BAG-domain proteins and E3s. Other UPS components such as deubiquitylating enzymes and PTMs of substrates or chaperones are also likely to play a role. In stress situations, the balance of these components is perturbed, which in turn affects the specificity of the PQC system and thus regulates the balance between folding and degradation. Accordingly, a system-wide understanding of the intricate proteostasis systems is necessary in order to predict the fate of a given misfolded protein (Powers and Gierasch, 2021). This point is also evident from our exploration of the importance of the yeast JDPs for the level of the RLLL degron. We note that multiple distinct JDPs affect the level of the RLLL-degron. Hence, most likely perturbation of the PQC system caused by deletion of the JDPs may also indirectly affect clearance of the RLLL degron. Although we observe protein interactions between the model substrate, Hsp70 and Ubr1, we cannot rule out that blocking Hsp70 does not indirectly affect Ubr1 and the UPS in general, and *vice versa.* However, an obvious advantage of Hsp70 and other chaperones playing a central role in PQC degradation is that PQC E3s in this way exploit a pre-existing cellular system capable of recognizing non-native proteins. From an evolutionary perspective, this seems intuitive, since the chaperone system arose early to regulate protein folding, whereas the UPS machinery is comparably new (Rebeaud et al., 2021).

**Figure 8.**
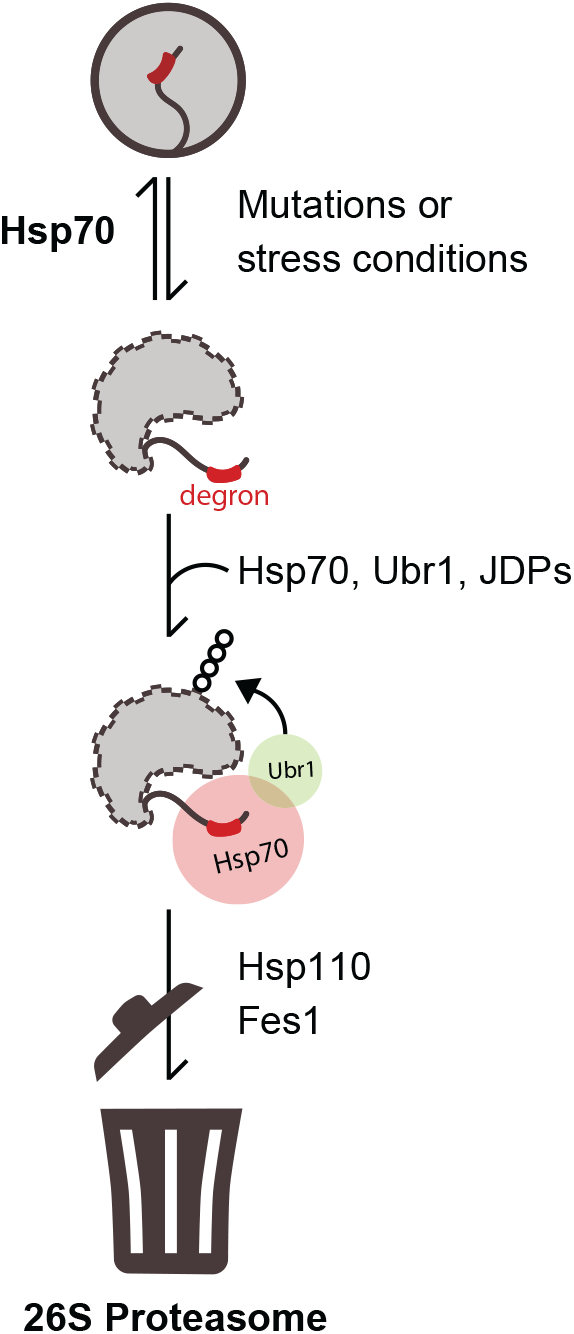
Schematic illustration of the proposed degradation pathway for misfolded proteins. Mutations or stress conditions increase the risk for protein misfolding leading to degron exposure. Hsp70 binds the degron motif in the misfolded protein, thus facilitating protein refolding (top) or degradation (bottom) in the presence of Ubr1 and JDPs. Ubr1-mediated ubiquitylation targets the protein for proteasomal degradation in collaboration with NEF proteins Hsp110 and Fes1.

A number of studies have aimed at clarifying the sequence features that constitute good Hsp70-binding elements (Clerico et al., 2021; Durme et al., 2009; McCarty et al., 1995; Nordquist et al., 2021; Rüdiger et al., 1997; Zhu et al., 1996). Degron sequences appear to hold similar features, and unsurprisingly, we find that residues flanking the degron motif also critically affect the degron activity. The sequence context likely determines the exposure of the degron but may also more directly affect degron recognition by chaperones and E3s. For instance, it has been shown that hydrophobic regions flanked by negative charges rarely engage with chaperones (Houben et al., 2020; Koopman and Rüdiger, 2020; Rousseau et al., 2006). This is supported by the observations presented here and by previous studies on cytosolic degrons (Johansson et al., 2022; Mashahreh et al., 2022; Maurer et al., 2016; Koren et al., 2018). We note, however, that efforts to define the discriminating features of PQC degrons are at least in part confounded by the overlapping nature of exposed hydrophobicity, the requirement for chaperone-binding, and insolubility.

Previous degron studies have also found high sequence variability and dependence on chaperones (Geffen et al., 2016; Maurer et al., 2016). Presumably, degron motifs must have a high sequence variability since the UPS must potentially be able to degrade the entire proteome. The genome thus encodes many different E3 ligases, each with different substrate preferences. However, since we find that Hsp70-binding motifs may act as degrons, the high sequence variability within degron motifs may also reflect the promiscuity of substrate binding to Hsp70 (Clerico et al., 2021; Rüdiger et al., 1997; Zhuravleva and Gierasch, 2015). Hence, unlike classical regulatory degrons such as the KEN-box, PQC degrons seem not to adhere to simple consensus motifs but rather to rely on the overall physical properties of the degron, such as hydrophobicity and charge patterns. This has also been shown for substrates of the yeast E3, San1, that targets misfolded nuclear proteins for degradation through direct interactions with exposed hydrophobic regions in the target protein (Gardner et al., 2005). Yet, even in the case of San1, chaperones can play a role both up- and downstream of the San1-catalyzed ubiquitylation (Guerriero et al., 2013; Jones et al., 2020; Kriegenburg et al., 2014).

In conclusion, the collective PQC degron catalogue may, to an extent, simply be sequence motifs that are recognized by PQC proteins including the abundant Hsp70. During stress, stable protein structures are destabilized and expose these sequences that take the role as degrons. Thus, a PQC degron can be any sequence or structure that is not normally found on a protein surface. Thus, much like the way the immune system only detects intruding antigens and not self-like motifs, the PQC detection system has evolved to recognize sequences that under normal conditions are not present (buried in native proteins). Accordingly, the presence of degron motifs in less structured proteins, like the intrinsically disordered proteins (IDPs), would necessarily be selected against, or in the case they are not, the degron would require activation, e.g. by phosphorylation or the protein would be expected to be highly unstable. Indeed this is what current studies suggest (Guharoy et al., 2016b) and reflected by our presented analyses of the yeast proteome.

## Materials and methods

### Yeast strains and plasmids

The BY4741 strain (*his3Δ1*, *leu2Δ0*, *met15Δ0*, *ura3Δ0*) of *S. cerevisiae* served as the wild-type control. Deletion strains were obtained from the Euroscarf collection. The *sse1-200sse2*Δ temperature-sensitive strain has been described before (Kaimal et al., 2017). Cells were grown in YPD (2% glucose, 2% peptone, 1% yeast extract) or synthetic complete (SC) media (2% glucose, 0.67% yeast nitrogen base without amino acids, 0.2% drop-out supplement) at 30 °C, unless otherwise stated. For high-level expression we used the pESC plasmid (Genscript) and glucose was substituted for galactose. The Ubr1-HA expression plasmid has been described before (Gowda et al., 2013). The pTR1412 plasmid for expressing the Ura3-HA-GFP fusion has been described before (Geffen et al., 2016). Transformation with DNA was performed as described previously (Gietz and Schiestl, 2007).

### Library construction and selection

Plasmids and polymerase chain reaction (PCR) products were purified using kits from Macherey-Nagel. PCR fragments were amplified using Q5 High-Fidelity 2x Master Mix (New England Biolabs). The primers (IDT) are listed in the Supplementary Table S1. The degron library was constructed using gap repair (Eckert-Boulet et al., 2012). The gap repair was performed with three PCR fragments amplified from and covering the entire pTR1412-1xRLLL plasmid, one of which contained a degron tag with 4 randomized codons across the RLLL motif (Trimer 20 codon mixes from IDT). Briefly, 0.1 pmol of fragment 1 (part of the *LEU2* marker, *URA3* and *GFP;* 4149 bp), 2 pmol of fragment 2 (degron tag 162 bp), and 0.1 pmol fragment 3 (missing part of the *LEU2* marker in fragment 1; 3828 bp) were transformed into BY4741 cells. Transformed colonies were scraped and pooled from 11 individual transformation plates. The estimated total number of colonies pooled was 120,000-130,000. 2 mL YPD was inoculated with the pooled library to reach OD_600nm_ = 0.3 and left to grow overnight at 30 °C. A 1:10 dilution of the culture was plated onto 0.8 mg/mL 5-FOA selection plates without leucine. Colonies resistant to 5-FOA were scraped off and pooled, and were estimated to represent <0.5% of the untreated library. ImageJ (version 1.51) was used for colony counting and set to include colonies between 10-1000 μm in size.

### Sequencing and statistical analyses

DNA was extracted from overnight cultures using Yeast DNA Extraction Kit (ThermoFisher). This served as a template in a PCR reaction amplifying 224 bp containing the randomized degron tag codons within 50 bp from one end. Purified PCR products from different samples were indexed using Nextera XT indexing. The indexed amplicons were quantified using Qubit 2.0 Fluorimeter (Life Technologies), pooled in a molar ratio of 1:3 for untreated (sample 1) and 5-FOA treated (sample 2) samples, respectively, and sequenced on Illumina MiSeq using 75-bp reads. The randomized degron tag codons were extracted from the sequencing reads using the cutadapt command-line tool (Martin, 2011) to remove flanking regions. All processed reads that did not have flanking regions removed or that after removal did not span exactly 12 nucleotides were discarded. Furthermore, we determined the number of reads needed for a trusted sequence to be 50 and 300 reads for samples 1 and 2, respectively by only allowing ~ 1 % sequence codons to be outside the set of 20 codons present in the Trimer 20 mix. In total, 18,000 and 883 different degron tag sequences, passed this cutoff from samples 1 and 2, respectively. Raw sequencing reads are available at doi: 10.17894/ucph.0f42dd43-a46c-4164-a80f-a7d3dd8f4a4d.

### Yeast growth assays

Cells in exponential growth phase were washed in sterile water and adjusted to OD_600nm_ = 0.4. A 1:5 serial dilution was spotted in 5 μL droplets on plates containing appropriate selection media and relevant chemical compounds: 1 mM bortezomib (LC laboratories), 0.8 mg/mL 5-FOA (Sigma). Plates were incubated at 30 °C, and growth phenotypes were analyzed after 48-72 hours.

### Microscopy

Expression was induced with 0.1 mM CuSO4 for 18 hours at 30 °C. Images were recorded using a Carl Zeiss Axio Vert.A1 microscope equipped with a 63x objective and a digital camera (Carl Zeiss AxioCam ICm1).

### SDS-PAGE and Western blotting

Protein levels in whole-cell lysates were analyzed by SDS-PAGE and Western blotting. Cycloheximide (Sigma) was used at a final concentration of 100 μg/mL. Bortezomib (LC Laboratories) was used at a final concentration of 1 mM. Myricetin (Sigma) was used at a final concentration of 100 μM. An equal number of cells (1-2×10^8^ cells) was harvested and lysed using glass beads and a FastPrep Bead Beating System (MP Biomedicals). For denaturing cell lysis, cells were lysed in 20% TCA (Sigma) and proteins were extracted by centrifugation and washed in 80% acetone. Finally, proteins were resuspended and boiled in 4x SDS sample buffer (250 mM Tris/HCl pH 6.8, 8% SDS, 40% glycerol, 0.05% bromophenol blue, 0.05% pyronin G, 2% β-mercaptoethanol). For analysis of protein solubility, proteins were lysed in 25 mM Tris/HCl pH 7.4, 50 mM NaCl, 10% glycerol and 2 mM MgCl2 and split into fractions by centrifugation (13,000 g, 30 min). Then, the volume of the pellet fractions was adjusted to match the volume of the supernatants prior to the addition of SDS sample buffer. The samples were separated by SDS-PAGE on 12.5% acrylamide gels and transferred to nitrocellulose membranes (Advantec). The membranes were blocked in 5% fat-free milk powder, 5 mM NaN3 in PBS (4.3 mM Na_2_HPO_4_, 1.47 mM KH_2_PO_4_, 137 mM NaCl, 2.7 mM KCl, pH 7.4). The primary antibodies and their sources were: anti-α-tubulin (Abcam, YL1/2 MA1-80017), anti-Pma1 (Abcam, 40B7 Ab4645), anti-GFP (Chromotek, 3H9 3h9-100), anti-Myc (Chromotek, 9E1 9e1-100), anti-HA (Roche, 15645900), anti-Pgk1 (Invitrogen, 22C5D8). Appropriate species-specific secondary antibodies conjugated to horseradish peroxidase were from DAKO.

### Co-immunoprecipitation

Cultures were incubated overnight in SC medium with 2% galactose instead of glucose to induce high-level expression from the pESC plasmid. Bortezomib (LC Laboratories) was added to a final concentration of 1 mM 4 hours prior to harvest. 1-2×10^8^ cells were lysed by mechanical lysis with glass beads in native lysis buffer (150 mM Tris/HCl pH 7.5, 50 mM NaCl, 1 mM PMSF and Complete protease inhibitors (Roche)). Cell debris was cleared from lysates by centrifugation (13,000 g, 30 min). The supernatants were incubated with GFP-trap resin (Chromotek) overnight at 4 °C. After extensive washing in lysis buffer, proteins were eluted with SDS sample buffer and analyzed by SDS-PAGE and Western blotting.

For the co-precipitation of Ubr1-HA with Ssa1-myc, we followed the above procedure but used a native lysis buffer containing 150 mM NaCl, 50 mM Hepes pH 7.4, 1 mM EDTA, 0.1% Triton X-100, 1 mM PMSF and Complete protease inhibitors (Roche), precleared cell extracts (10,000, 5 min) and incubated the supernatants with Myc-trap resin (Chromotek) for 3 hours at 4 °C.

### Circular dichroism

Amidated and acetylated peptides were synthesized by TAG Copenhagen A/S (Copenhagen, Denmark) with the following sequences: APPY-RLLL (CALLQSRLLLSAPRRAAATARY), APPY-DAAA (CALLQSDAAASAPRRAAATARY), APPY-var1 (CAAAQSRLLLSAPDDAAATARY), APPY-var7 (ATQLCSRLLLAYRRPAASALAR) and APPY-rev (YRATAAARRPASLLLRSQLLAC). Approximately 0.5 mg of each lyophilized peptide was weighed out and dissolved in 0.5 mL 10 mM sodium phosphate pH 7.0. After centrifugation at 15,000 g, 4 °C for 20 min, the supernatant was transferred to a new tube, and the peptide concentration was estimated from the absorbance at 280 nm and 214 nm (Kuipers and Gruppen, 2007). The peptides were diluted to 0.075 mg/mL (~30 μM) with 10 mM sodium phosphate pH 7.0, TCEP was added to a final concentration of 0.6 mM, and the pH was corrected to 7.0. A Jasco J-815 CD spectropolarimeter was used for recording Far-UV CD spectra from 260 nm to 190 nm at 20 °C with 1 nm bandwidth, 0.1 nm data pitch, 2 second D.I.T. and 10 nm/min. A total of 10 scans were accumulated and averaged. Similar spectra of the buffer alone were recorded with identical settings and subtracted.

### Limbo calculations

We used a local installation of the Limbo software (Durme et al., 2009) to predict Hsp70 binding for all peptides in the Ura3-HA-GFP-reporter library. Specifically, we performed Limbo calculations for each tetrapeptide in the context of the full 22 residue (APPY-like) peptide and used the strongest binding site predicted within each sequence as the value for that peptide. For the calculations on the yeast proteome, we calculated un-weighted Limbo scores of 7-residue fragments and assigned the central residue position using Limbo (Durme et al., 2009). The maximum score across 15 residues was subsequently assigned the central residue to obtain a score for a stretch totaling 22 residues, the length of the APPY peptide. We used yeast proteome abundance data collected by Dubreuil *et al.* (Dubreuil et al., 2019), and complemented these with half-life data for 1993 of the 4533 proteins (Christiano et al., 2014). As for abundance, proteins were assigned to one of 10 bins based on half-life with the same number of proteins in each bin. Proteins with any membrane-associated residues were excluded from this analysis to avoid that the generally hydrophobic patches give false positives. Limbo scores were assigned to each residue and the 30% highest scoring residues were considered Hsp70-binding sites. As a proxy for solvent exposure, the 30% of residues with the highest disorder (IUpred) score (Mészáros et al., 2018), as assigned by Dubreuil *et al.,* were considered disordered. The data is available at https://github.com/KULL-Centre/papers/tree/main/2021/Hsp70-Abildgaard-et-al.

### Mammalian cell growth and maintenance

Human HEK293T landing pad cells (TetBxb1BFPiCasp9 Clone 12) (Matreyek et al., 2020) were grown in Dulbecco’s Modified Eagle’s Medium (DMEM) with 0.32 mg/mL L-glutamine (Sigma Aldrich), 0.29 mg/mL Penicillin G potassium salt (BioChemica), 0.24 mg/mL Streptomycin sulphate (BioChemica), 10% (v/v) fetal bovine serum (FBS) (Sigma Aldrich) and 2 μg/mL doxycycline (Sigma-Aldrich). Cells were passaged when they were at 70-80% confluency and were detached with trypsin 0.25% (Difco). The cell lines used in this study tested negative for mycoplasma.

### Integration of APPY and APPY variants constructs into HEK293T landing pad cells

The APPY and APPY variant constructs were integrated into the HEK293T landing pad cells by transfecting 1.2 μg of the APPY recombination plasmid along with 0.08 μg of the integrase vector pCAG-NLS-Bxb1 (17,5:1 molar ratio). The transfection reagent was Fugene HD (Promega). Two days after the transfection recombinant cells were selected by adding 2 μg/mL of doxycycline (dox) and 10 nM of AP1903 (MedChemExpress). Cells were selected for 48 hours after which they were allowed to grow for another 48 hours to ensure the expression of the transgenes reached steady state levels before performing flow cytometry profiling.

### HEK293T drug treatments and flow cytometry profiling

For the YM01 treatment cells were treated with 5 μM of YM01 (StressMarq Biosciences) for 22 hours before flow-cytometry profiling. For bortezomib treatment cells were treated with 15 μM of bortezomib (LC Laboratories) for 5 hours before flow-cytometry profiling. On the day of flowcytometry profiling, cells were washed with PBS, detached with trypsin, and resuspended in PBS with 2% (v/v) FBS. The suspension of cells was filtered through a 50 μm mesh filter and transferred to a 5 mL tube. Samples were analyzed by FACSJazz (BD) or FACS Aria III (BD) (bortezomib experiment). Single cells were gated by using forward and side scatter. Then recombinant cells were gated based on their lack of BFP signal and presence of mCherry signal. Finally, the GFP:mCherry ratios were quantified. The laser used for excitation of BFP was 405 nm, while for GFP it was 488 nm, and for mCherry 561 nm with FACSJazz and 562 nm with FACS AriaIII. The filters used were 450/50 for BFP, 530/40 for GFP and 610/20 for mCherry for FACSJazz and 442/46 for BFP, 530/30 for GFP and 615/20 for mCherry for the FACS Aria III.

### Reverse transcription quantitative real-time PCR

HSPA1A and HSPA1B (HSPA1A/B) mRNA levels were quantified using mRNA from 0.5-2×10^6^ cells that were harvested 9 days after the addition of dox. The mRNA was extracted using the RNeasy mini kit (Qiagen). After eliminating possible DNA contamination by DNase I (Invitrogen) treatment, reverse transcription was performed for 1 μg of mRNA by using oligo-dTs and the Maxima H Minus Reverse Transcriptase (Thermo Scientific). Approximately 50 ng of cDNA were used to perform realtime qPCR with the Maxima SYBR Green/ROX qPCR Master Mix (Thermo Scientific) according to the manufacturer’s instructions. The primers used for HSPA1A/B cDNA and the housekeeping ACTB cDNA amplifications are listed in the supplemental material (Table S1).

## Acknowledgements

The authors thank Anne-Marie Lauridsen and Søren Lindemose for excellent technical assistance, Peter S. Millard for help with the circular dichroism, Asta B. Andersen for helping with the illustrations, and Frederikke V. Larsen for assistance early in the project. We thank Profs. F. Rousseau and J. Schymkowitz (VIB) for sharing the Limbo Software.

## Conflicts of interest

No conflicting interests to declare.

## Funding

This work was supported by the Novo Nordisk Foundation (https://novonordiskfonden.dk) challenge programme PRISM (NNF18OC0033950; to A.S, K.L.L. & R.H.P.), REPIN (NNF18OC0033926; to R.H.P.) and NNF18OC0052441 and NNF0071057 (to R.H.P.), the Lundbeck Foundation (https://www.lundbeckfonden.com) R249-2017-510 (to R.H.P.) and R272-2017-4528 (to A.S.), the Danish Council for Independent Research (Natur og Univers, Det Frie Forskningsråd) (https://dff.dk/) 7014-00039B (to R.H.P.), the Villum Foundation (https://veluxfoundations.dk/) 40526 (to R.H.P.), and the European Commission Horizon 2020 programme (MIAMi: No. 722287) (to M.K.J.). The funders had no role in study design, data collection and analysis, decision to publish, or preparation of the manuscript.

## Author contributions

A.B.A., V.V., S.D.P., F.B.L., C.K. and K.E.J. conducted the experiments. Data analyses by A.B.A., V.V., S.D.P, K.E.J., A.S., T.R., C.A., M.K.J., K.L.L., and R.H.P. Experimental design by A.B.A., S.D.P., M.K.J., T.R., and R.H.P. A.B.A., C.A., K.L.L., and R.H.P. conceived the study. A.B.A. and R.H.P. wrote the paper.

## Notes

### Competing Interest Statement

The authors have declared no competing interest.

### Summary of Updates

Some changes to the text and figures.

## References

Abildgaard, A.B., Gersing, S.K., Larsen-Ledet, S., Nielsen, S.V., Stein, A., Lindorff-Larsen, K., Hartmann-Petersen, R., 2020. Co-Chaperones in Targeting and Delivery of Misfolded Proteins to the 26S Proteasome. Biomolecules 10. https://doi.org/10.3390/biom10081141

Abildgaard, A.B., Stein, A., Nielsen, S.V., Schultz-Knudsen, K., Papaleo, E., Shrikhande, A., Hoffmann, E.R., Bernstein, I., Gerdes, A.-M., Takahashi, M., Ishioka, C., Lindorff-Larsen, K., Hartmann-Petersen, R., 2019. Computational and cellular studies reveal structural destabilization and degradation of MLH1 variants in Lynch syndrome. eLife 8, e49138. https://doi.org/10.7554/eLife.49138

Amm, I., Norell, D., Wolf, D.H., 2015. Absence of the Yeast Hsp31 Chaperones of the DJ-1 Superfamily Perturbs Cytoplasmic Protein Quality Control in Late Growth Phase. PLoS One 10, e0140363. https://doi.org/10.1371/journal.pone.0140363

Andréasson, C., Fiaux, J., Rampelt, H., Mayer, M.P., Bukau, B., 2008. Hsp110 Is a Nucleotide-activated Exchange Factor for Hsp70. J. Biol. Chem. 283, 8877–8884. https://doi.org/10.1074/jbc.M710063200

Bachmair, A., Finley, D., Varshavsky, A., 1986. In vivo half-life of a protein is a function of its amino-terminal residue. Science 234, 179–186. https://doi.org/10.1126/science.3018930

Balchin, D., Hayer-Hartl, M., Hartl, F.U., 2020. Recent advances in understanding catalysis of protein folding by molecular chaperones. FEBS Letters 594, 2770–2781. https://doi.org/10.1002/1873-3468.13844

Balchin, D., Hayer-Hartl, M., Hartl, F.U., 2016. In vivo aspects of protein folding and quality control. Science 353. https://doi.org/10.1126/science.aac4354

Bates, G.P., Dorsey, R., Gusella, J.F., Hayden, M.R., Kay, C., Leavitt, B.R., Nance, M., Ross, C.A., Scahill, R.I., Wetzel, R., Wild, E.J., Tabrizi, S.J., 2015. Huntington disease. Nat Rev Dis Primers 1, 15005. https://doi.org/10.1038/nrdp.2015.5

Boeke, J.D., LaCroute, F., Fink, G.R., 1984. A positive selection for mutants lacking orotidine-5’-phosphate decarboxylase activity in yeast: 5-fluoro-orotic acid resistance. Mol Gen Genet 197, 345–346. https://doi.org/10.1007/BF00330984

Bucciantini, M., Giannoni, E., Chiti, F., Baroni, F., Formigli, L., Zurdo, J., Taddei, N., Ramponi, G., Dobson, C.M., Stefani, M., 2002. Inherent toxicity of aggregates implies a common mechanism for protein misfolding diseases. Nature 416, 507–511. https://doi.org/10.1038/416507a

Chang, L., Miyata, Y., Ung, P.M.U., Bertelsen, E.B., McQuade, T.J., Carlson, H.A., Zuiderweg, E.R.P., Gestwicki, J.E., 2011. Chemical Screens against a Reconstituted Multiprotein Complex: Myricetin Blocks DnaJ Regulation of DnaK through an Allosteric Mechanism. Chemistry & Biology 18, 210–221. https://doi.org/10.1016/j.chembiol.2010.12.010

Christiano, R., Nagaraj, N., Fröhlich, F., Walther, T.C., 2014. Global Proteome Turnover Analyses of the Yeasts S. cerevisiae and S. pombe. Cell Reports 9, 1959–1965. https://doi.org/10.1016/j.celrep.2014.10.065

Clerico, E.M., Pozhidaeva, A.K., Jansen, R.M., Özden, C., Tilitsky, J.M., Gierasch, L.M., 2021. Selective promiscuity in the binding of E. coli Hsp70 to an unfolded protein. PNAS 118. https://doi.org/10.1073/pnas.2016962118

Dahiya, V., Buchner, J., 2019. Functional principles and regulation of molecular chaperones. Adv Protein Chem Struct Biol 114, 1–60. https://doi.org/10.1016/bs.apcsb.2018.10.001

Dikic, I., 2017. Proteasomal and Autophagic Degradation Systems. Annu. Rev. Biochem. 86, 193–224. https://doi.org/10.1146/annurev-biochem-061516-044908

Dubreuil, B., Matalon, O., Levy, E.D., 2019. Protein Abundance Biases the Amino Acid Composition of Disordered Regions to Minimize Non-functional Interactions. Journal of Molecular Biology 431, 4978–4992. https://doi.org/10.1016/j.jmb.2019.08.008

Durme, J.V., Maurer-Stroh, S., Gallardo, R., Wilkinson, H., Rousseau, F., Schymkowitz, J., 2009. Accurate Prediction of DnaK-Peptide Binding via Homology Modelling and Experimental Data. PLOS Computational Biology 5, e1000475. https://doi.org/10.1371/journal.pcbi.1000475

Eckert-Boulet, N., Pedersen, M.L., Krogh, B.O., Lisby, M., 2012. Optimization of ordered plasmid assembly by gap repair in Saccharomyces cerevisiae. Yeast 29, 323–334. https://doi.org/10.1002/yea.2912

Fredrickson, E.K., Gallagher, P.S., Clowes Candadai, S.V., Gardner, R.G., 2013. Substrate recognition in nuclear protein quality control degradation is governed by exposed hydrophobicity that correlates with aggregation and insolubility. J Biol Chem 288, 6130–6139. https://doi.org/10.1074/jbc.M112.406710

Gallagher, P.S., Clowes Candadai, S.V., Gardner, R.G., 2014. The requirement for Cdc48/p97 in nuclear protein quality control degradation depends on the substrate and correlates with substrate insolubility. Journal of Cell Science 127, 1980–1991. https://doi.org/10.1242/jcs.141838

Gardner, R.G., Nelson, Z.W., Gottschling, D.E., 2005. Degradation-mediated protein quality control in the nucleus. Cell 120, 803–815. https://doi.org/10.1016/j.cell.2005.01.016

Geffen, Y., Appleboim, A., Gardner, R.G., Friedman, N., Sadeh, R., Ravid, T., 2016. Mapping the Landscape of a Eukaryotic Degronome. Mol. Cell 63, 1055–1065. https://doi.org/10.1016/j.molcel.2016.08.005

Gersing, S.K., Wang, Y., Grønbæk-Thygesen, M., Kampmeyer, C., Clausen, L., Willemoës, M., Andréasson, C., Stein, A., Lindorff-Larsen, K., Hartmann-Petersen, R., 2021. Mapping the degradation pathway of a disease-linked aspartoacylase variant. PLOS Genetics 17, e1009539. https://doi.org/10.1371/journal.pgen.1009539

Gietz, R.D., Schiestl, R.H., 2007. High-efficiency yeast transformation using the LiAc/SS carrier DNA/PEG method. Nat Protoc 2, 31–34. https://doi.org/10.1038/nprot.2007.13

Gilon, T., Chomsky, O., Kulka, R.G., 1998. Degradation signals for ubiquitin system proteolysis in Saccharomyces cerevisiae. EMBO J 17, 2759–2766. https://doi.org/10.1093/emboj/17.10.2759

Gowda, N.K.C., Kaimal, J.M., Kityk, R., Daniel, C., Liebau, J., Öhman, M., Mayer, M.P., Andréasson, C., 2018. Nucleotide exchange factors Fes1 and HspBP1 mimic substrate to release misfolded proteins from Hsp70. Nat. Struct. Mol. Biol. 25, 83–89. https://doi.org/10.1038/s41594-017-0008-2

Gowda, N.K.C., Kandasamy, G., Froehlich, M.S., Dohmen, R.J., Andréasson, C., 2013. Hsp70 nucleotide exchange factor Fes1 is essential for ubiquitin-dependent degradation of misfolded cytosolic proteins. PNAS 110, 5975–5980. https://doi.org/10.1073/pnas.1216778110

Guerriero, C.J., Weiberth, K.F., Brodsky, J.L., 2013. Hsp70 targets a cytoplasmic quality control substrate to the San1p ubiquitin ligase. J. Biol. Chem. 288, 18506–18520. https://doi.org/10.1074/jbc.M113.475905

Guharoy, M., Bhowmick, P., Sallam, M., Tompa, P., 2016a. Tripartite degrons confer diversity and specificity on regulated protein degradation in the ubiquitin-proteasome system. Nat Commun 7, 10239. https://doi.org/10.1038/ncomms10239

Guharoy, M., Bhowmick, P., Tompa, P., 2016b. Design Principles Involving Protein Disorder Facilitate Specific Substrate Selection and Degradation by the Ubiquitin-Proteasome System. J Biol Chem 291, 6723–6731. https://doi.org/10.1074/jbc.R115.692665

Heck, J.W., Cheung, S.K., Hampton, R.Y., 2010. Cytoplasmic protein quality control degradation mediated by parallel actions of the E3 ubiquitin ligases Ubr1 and San1. Proc. Natl. Acad. Sci. U.S.A. 107, 1106–1111. https://doi.org/10.1073/pnas.0910591107

Hickey, C.M., Breckel, C., Zhang, M., Theune, W.C., Hochstrasser, M., 2021. Protein quality control degron-containing substrates are differentially targeted in the cytoplasm and nucleus by ubiquitin ligases. Genetics 217, 1–19. https://doi.org/10.1093/genetics/iyaa031

Höhfeld, J., Jentsch, S., 1997. GrpE-like regulation of the Hsc70 chaperone by the anti-apoptotic protein BAG-1. The EMBO Journal 16, 6209–6216. https://doi.org/10.1093/emboj/16.20.6209

Houben, B., Michiels, E., Ramakers, M., Konstantoulea, K., Louros, N., Verniers, J., van der Kant, R., De Vleeschouwer, M., Chicória, N., Vanpoucke, T., Gallardo, R., Schymkowitz, J., Rousseau, F., 2020. Autonomous aggregation suppression by acidic residues explains why chaperones favour basic residues. EMBO J. e102864. https://doi.org/10.15252/embj.2019102864

Johansson, K.E., Mashahreh, B., Hartmann-Petersen, R., Ravid, T., Lindorff-Larsen, K., 2022. Prediction of quality-control degradation signals in yeast proteins. https://doi.org/10.1101/2022.04.06.487301

Johnson, P.R., Swanson, R., Rakhilina, L., Hochstrasser, M., 1998. Degradation Signal Masking by Heterodimerization of MATα2 and MATa1 Blocks Their Mutual Destruction by the Ubiquitin-Proteasome Pathway. Cell 94, 217–227. https://doi.org/10.1016/S0092-8674(00)81421-X

Jones, R.D., Enam, C., Ibarra, R., Borror, H.R., Mostoller, K.E., Fredrickson, E.K., Lin, J., Chuang, E., March, Z., Shorter, J., Ravid, T., Kleiger, G., Gardner, R.G., 2020. The extent of Ssa1/Ssa2 Hsp70 chaperone involvement in nuclear protein quality control degradation varies with the substrate. Mol. Biol. Cell 31, 221–233. https://doi.org/10.1091/mbc.E18-02-0121

Kaimal, J.M., Kandasamy, G., Gasser, F., Andréasson, C., 2017. Coordinated Hsp110 and Hsp104 Activities Power Protein Disaggregation in Saccharomyces cerevisiae. Mol Cell Biol 37, e00027–17. https://doi.org/10.1128/MCB.00027-17

Kampmeyer, C., Larsen-Ledet, S., Wagnkilde, M.R., Michelsen, M., Iversen, H.K.M., Nielsen, S.V., Lindemose, S., Caregnato, A., Ravid, T., Stein, A., Teilum, K., Lindorff-Larsen, K., Hartmann-Petersen, R., 2022. Disease-linked mutations cause exposure of a protein quality control degron. Structure S0969-2126(22)00190-3. https://doi.org/10.1016/j.str.2022.05.016

Kandasamy, G., Andréasson, C., 2018. Hsp70-Hsp110 chaperones deliver ubiquitin-dependent and - independent substrates to the 26S proteasome for proteolysis in yeast. J. Cell. Sci. 131. https://doi.org/10.1242/jcs.210948

Kohler, V., Andréasson, C., 2020. Hsp70-mediated quality control: should I stay or should I go? Biol Chem 401, 1233–1248. https://doi.org/10.1515/hsz-2020-0187

Koopman, M.B., Rüdiger, S.G., 2020. Behind closed gates - chaperones and charged residues determine protein fate. EMBO J 39, e104939. https://doi.org/10.15252/embj.2020104939

Koren, I., Timms, R.T., Kula, T., Xu, Q., Li, M.Z., Elledge, S.J., 2018. The Eukaryotic Proteome Is Shaped by E3 Ubiquitin Ligases Targeting C-Terminal Degrons. Cell 173, 1622–1635.e14. https://doi.org/10.1016/j.cell.2018.04.028

Kriegenburg, F., Jakopec, V., Poulsen, E.G., Nielsen, S.V., Roguev, A., Krogan, N., Gordon, C., Fleig, U., Hartmann-Petersen, R., 2014. A Chaperone-Assisted Degradation Pathway Targets Kinetochore Proteins to Ensure Genome Stability. PLOS Genetics 10, e1004140. https://doi.org/10.1371/journal.pgen.1004140

Kuipers, B.J.H., Gruppen, H., 2007. Prediction of Molar Extinction Coefficients of Proteins and Peptides Using UV Absorption of the Constituent Amino Acids at 214 nm To Enable Quantitative Reverse Phase High-Performance Liquid Chromatography-Mass Spectrometry Analysis. J. Agric. Food Chem. 55, 5445–5451. https://doi.org/10.1021/jf070337l

Kwon, Y.T., Ciechanover, A., 2017. The Ubiquitin Code in the Ubiquitin-Proteasome System and Autophagy. Trends Biochem Sci 42, 873–886. https://doi.org/10.1016/j.tibs.2017.09.002

Macossay-Castillo, M., Marvelli, G., Guharoy, M., Jain, A., Kihara, D., Tompa, P., Wodak, S.J., 2019. The Balancing Act of Intrinsically Disordered Proteins: Enabling Functional Diversity while Minimizing Promiscuity. Journal of Molecular Biology 431, 1650–1670. https://doi.org/10.1016/j.jmb.2019.03.008

Martin, M., 2011. Cutadapt removes adapter sequences from high-throughput sequencing reads. EMBnet.journal 17, 10–12. https://doi.org/10.14806/ej.17.1.200

Mashahreh, B., Armony, S., Johansson, K.E., Chapelbaum, A., Friedman, N., Gardner, R.G., Hartmann-Petersen, R., Lindorff-Larsen, K., Ravid, T., 2022. Conserved degronome features governing quality control-associated proteolysis. https://doi.org/10.1101/2022.04.06.487275

Matreyek, K.A., Stephany, J.J., Chiasson, M.A., Hasle, N., Fowler, D.M., 2020. An improved platform for functional assessment of large protein libraries in mammalian cells. Nucleic Acids Res 48, e1. https://doi.org/10.1093/nar/gkz910

Maurer, M.J., Spear, E.D., Yu, A.T., Lee, E.J., Shahzad, S., Michaelis, S., 2016. Degradation Signals for Ubiquitin-Proteasome Dependent Cytosolic Protein Quality Control (CytoQC) in Yeast. G3 (Bethesda) 6, 1853–1866. https://doi.org/10.1534/g3.116.027953

McCarty, J.S., Buchberger, A., Reinstein, J., Bukau, B., 1995. The role of ATP in the functional cycle of the DnaK chaperone system. J. Mol. Biol. 249, 126–137. https://doi.org/10.1006/jmbi.1995.0284

Mészáros, B., Erdos, G., Dosztányi, Z., 2018. IUPred2A: context-dependent prediction of protein disorder as a function of redox state and protein binding. Nucleic Acids Res 46, W329–W337. https://doi.org/10.1093/nar/gky384

Montgomery, D.L., Morimoto, R.I., Gierasch, L.M., 1999. Mutations in the substrate binding domain of the Escherichia coli 70 kda molecular chaperone, DnaK, which alter substrate affinity or interdomain coupling11 Edited by M. Gottesman. Journal of Molecular Biology 286, 915–932. https://doi.org/10.1006/jmbi.1998.2514

Nordquist, E.B., English, C.A., Clerico, E.M., Sherman, W., Gierasch, L.M., Chen, J., 2021. Physics-based modeling provides predictive understanding of selectively promiscuous substrate binding by Hsp70 chaperones. PLOS Computational Biology 17, e1009567. https://doi.org/10.1371/journal.pcbi.1009567

Oh, E., Akopian, D., Rape, M., 2018. Principles of Ubiquitin-Dependent Signaling. Annu. Rev. Cell Dev. Biol. 34, 137–162. https://doi.org/10.1146/annurev-cellbio-100617-062802

Pfleger, C.M., Kirschner, M.W., 2000. The KEN box: an APC recognition signal distinct from the D box targeted by Cdh1. Genes Dev 14, 655–665.

Poewe, W., Seppi, K., Tanner, C.M., Halliday, G.M., Brundin, P., Volkmann, J., Schrag, A.-E., Lang, A.E., 2017. Parkinson disease. Nat Rev Dis Primers 3, 1–21. https://doi.org/10.1038/nrdp.2017.13

Pohl, C., Dikic, I., 2019. Cellular quality control by the ubiquitin-proteasome system and autophagy. Science 366, 818–822. https://doi.org/10.1126/science.aax3769

Powers, E.T., Gierasch, L.M., 2021. The Proteome Folding Problem and Cellular Proteostasis. J Mol Biol 167197. https://doi.org/10.1016/j.jmb.2021.167197

Ravid, T., Kreft, S.G., Hochstrasser, M., 2006. Membrane and soluble substrates of the Doa10 ubiquitin ligase are degraded by distinct pathways. EMBO J 25, 533–543. https://doi.org/10.1038/sj.emboj.7600946

Rebeaud, M.E., Mallik, S., Goloubinoff, P., Tawfik, D.S., 2021. On the evolution of chaperones and cochaperones and the expansion of proteomes across the Tree of Life. Proc Natl Acad Sci U S A 118, e2020885118. https://doi.org/10.1073/pnas.2020885118

Rosenzweig, R., Nillegoda, N.B., Mayer, M.P., Bukau, B., 2019. The Hsp70 chaperone network. Nat. Rev. Mol. Cell Biol. 20, 665–680. https://doi.org/10.1038/s41580-019-0133-3

Rousseau, F., Serrano, L., Schymkowitz, J.W.H., 2006. How Evolutionary Pressure Against Protein Aggregation Shaped Chaperone Specificity. Journal of Molecular Biology 355, 1037–1047. https://doi.org/10.1016/j.jmb.2005.11.035

Rüdiger, S., Germeroth, L., Schneider-Mergener, J., Bukau, B., 1997. Substrate specificity of the DnaK chaperone determined by screening cellulose-bound peptide libraries. EMBO J. 16, 1501–1507. https://doi.org/10.1093/emboj/16.7.1501

Samant, R.S., Livingston, C.M., Sontag, E.M., Frydman, J., 2018. Distinct proteostasis circuits cooperate in nuclear and cytoplasmic protein quality control. Nature 563, 407–411. https://doi.org/10.1038/s41586-018-0678-x

Schmid, D., Baici, A., Gehring, H., Christen, P., 1994. Kinetics of molecular chaperone action. Science 263, 971–973. https://doi.org/10.1126/science.8310296

Schmid, D., Jaussi, R., Christen, P., 1992. Precursor of mitochondrial aspartate aminotransferase synthesized in Escherichia coli is complexed with heat-shock protein DnaK. Eur J Biochem 208, 699–704. https://doi.org/10.1111/j.1432-1033.1992.tb17237.x

Shiber, A., Breuer, W., Brandeis, M., Ravid, T., 2013. Ubiquitin conjugation triggers misfolded protein sequestration into quality control foci when Hsp70 chaperone levels are limiting. Mol Biol Cell 24, 2076–2087. https://doi.org/10.1091/mbc.E13-01-0010

Singh, A., Vashistha, N., Heck, J., Tang, X., Wipf, P., Brodsky, J.L., Hampton, R.Y., 2020. Direct involvement of Hsp70 ATP hydrolysis in Ubr1-dependent quality control. MBoC 31, 2669–2686. https://doi.org/10.1091/mbc.E20-08-0541

Summers, D.W., Wolfe, K.J., Ren, H.Y., Cyr, D.M., 2013. The Type II Hsp40 Sis1 Cooperates with Hsp70 and the E3 Ligase Ubr1 to Promote Degradation of Terminally Misfolded Cytosolic Protein. PLoS One 8, e52099. https://doi.org/10.1371/journal.pone.0052099

Timms, R.T., Koren, I., 2020. Tying up loose ends: the N-degron and C-degron pathways of protein degradation. Biochem Soc Trans 48, 1557–1567. https://doi.org/10.1042/BST20191094

Varshavsky, A., 2011. The N-end rule pathway and regulation by proteolysis. Protein Sci 20, 1298–1345. https://doi.org/10.1002/pro.666

Varshavsky, A., 1991. Naming a targeting signal. Cell 64, 13–15. https://doi.org/10.1016/0092-8674(91)90202-a

Zhang, P., Leu, J.I.-J., Murphy, M.E., George, D.L., Marmorstein, R., 2014. Crystal structure of the stress-inducible human heat shock protein 70 substrate-binding domain in complex with peptide substrate. PLoS One 9, e103518. https://doi.org/10.1371/journal.pone.0103518

Zhu, X., Zhao, X., Burkholder, W.F., Gragerov, A., Ogata, C.M., Gottesman, M.E., Hendrickson, W.A., 1996. Structural analysis of substrate binding by the molecular chaperone DnaK. Science 272, 1606–1614. https://doi.org/10.1126/science.272.5268.1606

Zhuravleva, A., Gierasch, L.M., 2015. Substrate-binding domain conformational dynamics mediate Hsp70 allostery. PNAS 112, E2865–E2873.

