## Supplemental Figures and Tables for "HSP70-binding motifs function as protein quality control degrons"

### Supplemental Material

**Supplemental Figure S1**, Effect of chaperones and co-chaperones p. 2
**Supplemental Figure S2**, Circular dichroism spectroscopy of selected peptides p. 3
**Supplemental Table S1**, DNA primers used in this study p. 4

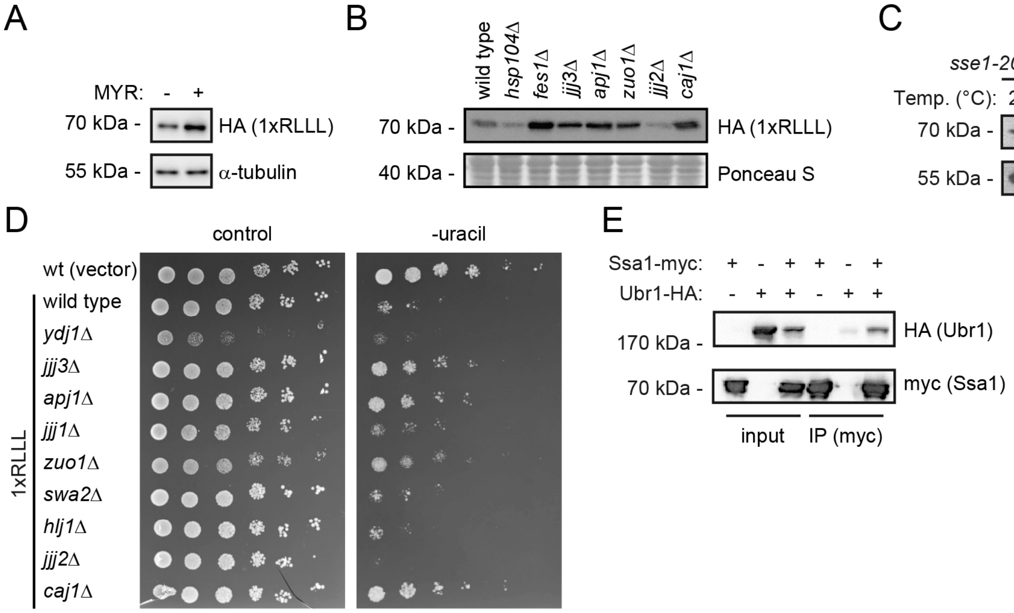


**Supplementary Figure S1** *Effect of chaperones and co-chaperones.* (A) The protein levels of the RLLL degron from cells treated (+) or untreated (-) with the Hsp70-inhibitor myricetin (MYR) for 16 hours were compared by SDS-PAGE and blotting against the HA-tag on the reporter. Tubulin served as loading control. (B) The protein levels of the RLLL degron were compared in the indicated yeast strains by SDS-PAGE and blotting for the HA-tag on the reporter. A Ponceau S staining of the membrane is included as loading control. (C) Protein levels in the temperature-sensitive (ts) Hsp110-double mutant strain (*sse1-200sse2Δ*) at 25 °C and 30 °C were compared by blotting against the HA-tag. Tubulin served as loading control. (D) The dependence of co-chaperones for targeting the RLLL degron was analyzed by growth assays on solid media using the indicated null mutants. Wild-type cells transformed with the reporter vector alone were included for comparison. (E) Wild-type yeast cells expressing myc-tagged Ssa1 and/or HA-tagged Ubr1, as indicated, were used for immunoprecipitation (IP) with myc-trap resin. The precipitated material was analyzed by SDS-PAGE and western blotting using antibodies to myc and HA.

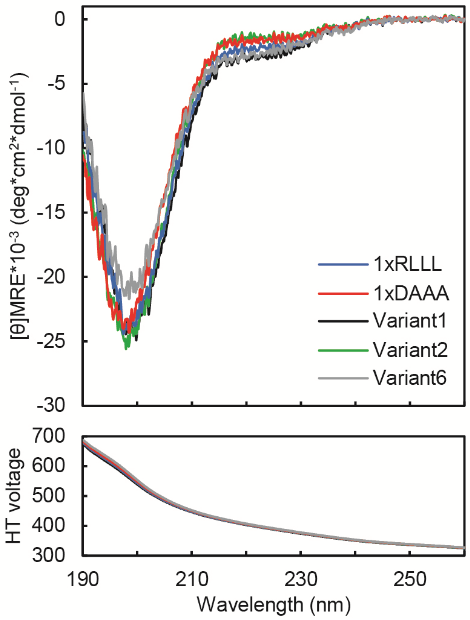


**Supplementary Figure S2** *Circular dichroism spectroscopy of selected peptides.* The secondary structure of selected variants was analyzed by far-UV circular dichroism (CD) spectroscopy. The top panel shows molar ellipticity and the lower panel shows HT voltage.

**Supplementary Table S1**

*DNA primers used in this study*

| Name | Sequence | Purpose |
|---|---|---|
| F1 _LEU2_1119_ fw | GCTGGTGATTATAATACCATT TAGGTGGGTTGG | Forward fragment 1 containing URA3-GFP for library construction |
| F1_degron_ta g_18 _rv | TGATTGTAACAATGCACAGG ATCCTTCGTC | Reverse fragment 1 containing URA3-GFP for library construction |
| F2_degron_ta g_fw | GACGAAGGATCCTGTGCATT GTTACAATCA /iTriMix20//iTriMix20//iTriMix20/ /iTriMix20/ TCAGCACCAAGAAGAGCTGC | Forward fragment 2 containing a degron tag including four trimer 20 codon mixes for library construction |
| F2_rv | ATTACGCCAAGCTCGAAATT AACCC | Reverse fragment 2 containing a degron tag for library construction |
| F3_fw | AGTGAGGGTTAATTTCGAGC TTGGC | Forward fragment 3 containing origin of replication for library construction |
| F3_LEU2_73 5_rv | CCAACCCACCTAAATGGTAT TATAATCACC | Reverse fragment 3 containing origin of replication for library construction |
| Illumina_adap ter _fw | TCGTCGGCAGCGTCAGATGT GTATAAGAGACAG cctacgatcgacgaagg | Forward containing degron tag for library sequencing |
| Illumina_adap ter _rv | GTCTCGTGGGCTCGGAGATG TGTATAAGAGACAG tgtggaattgtgagcggata | Reverse containing degron tag for library sequencing |
| VV48 | GGGTGCTGATCCAGGTGTAC | qPCR HSPA1A/B forward |
| VV49 | GTCGAAGGTCACCTCGATCT G | qPCR HSPA1A/B reverse |
| VV52 | GGCCAGGTCATCACCATTGG | qPCR ACTB forward |
| VV53 | CGGATGTCCACGTCACACTTC | qPCR ACTB reverse |
